# The genome of the sapphire damselfish *Chrysiptera cyanea*: a new resource to support further investigation of the evolution of Pomacentrids

**DOI:** 10.1101/2024.11.06.622371

**Authors:** Emma Gairin, Saori Miura, Hiroki Takamiyagi, Marcela Herrera, Vincent Laudet

**Author notes:** Co last authors. Corresponding author: Emma Gairin.

## Abstract

The number of high-quality genomes is rapidly growing across taxa. However, it remains limited for coral reef fish of the Pomacentrid family, with most research focused on anemonefish. Here, we present the first assembly for a Pomacentrid of the genus *Chrysiptera*. Using PacBio long-read sequencing with a coverage of 94.5x, the genome of the Sapphire Devil, *Chrysiptera cyanea* was assembled and annotated. The final assembly consisted of 896 Mb pairs across 91 contigs, with a BUSCO completeness of 97.6%. 28,173 genes were identified. Comparative analyses with available chromosome-scale assemblies for related species identified contig-chromosome correspondences. This genome will be useful to use as a comparison to study the specific adaptations linked to symbiosis life of the closely related anemonefish. Furthermore, this species is present in most tropical coastal areas in the Indo-West Pacific and could become a model for environmental monitoring. This work will allow to expand coral reef research efforts and highlights the power of long-read assemblies to retrieve high quality genomes.

## Introduction

Damselfish (Pomacentridae family) are highly abundant and common demersal reef fish throughout temperate, subtropical, and tropical waters. Many damselfish species, such as anemonefish, are prominent on the aquariology trade worldwide. They are also commonly studied to answer a number of scientific questions, notably at the ecological, behavioural, and developmental levels. As a key resource to deepen scientific analyses, genomes have been generated for fifteen species of Pomacentrids, including ten *Amphiprion* species [1–7]. Chromosome-level genomes are currently available for five species: four Pomacentrinae (*Acanthochromis polyacanthus* [8]*, Amphiprion ocellaris* [4]*, Amphiprion percula* [1]*, Amphiprion clarkii* [9], and one Chrominae, *Dascyllus trimaculatus* [3]. To expand the genomic resources of Pomacentrids further and reduce the bias for *Amphiprion,* we present here the first genome for the Cheiloprionini *Chrysiptera cyanea*, a member of the Pomacentrinae subfamily.

The Sapphire Devil *Chrysiptera cyanea* (Quoy and Gaimard 1825, p392 [10]) is a strikingly blue damselfish (Figure 1A) that can be encountered on shallow (0-10 meters) Indo-West Pacific tropical coral reefs [11] (Figure 1C). *C. cyanea* is territorial, reef-associated, and non-migratory. It is most generally found around rubble and corals in subtidal reef flats and tide pools (Figure 1B). *C. cyanea* has an omnivorous diet, commonly consisting of plankton, algae, small benthic crustaceans. The size of females typically ranges between 38 and 54 mm TL (total length), while males measure 49 to 73 mm TL [12]. *C. cyanea* live in small to large schools that typically consist of one or a few males and more females [13,14] (Figure 1B). The reproductive season of *Chrysiptera cyanea* in Okinawa lasts from April to August [15,16], and large densities of juvenile fish can be found in clusters along the coastline during the reproduction season in the presence or not of adult conspecifics.

**Figure 1.**
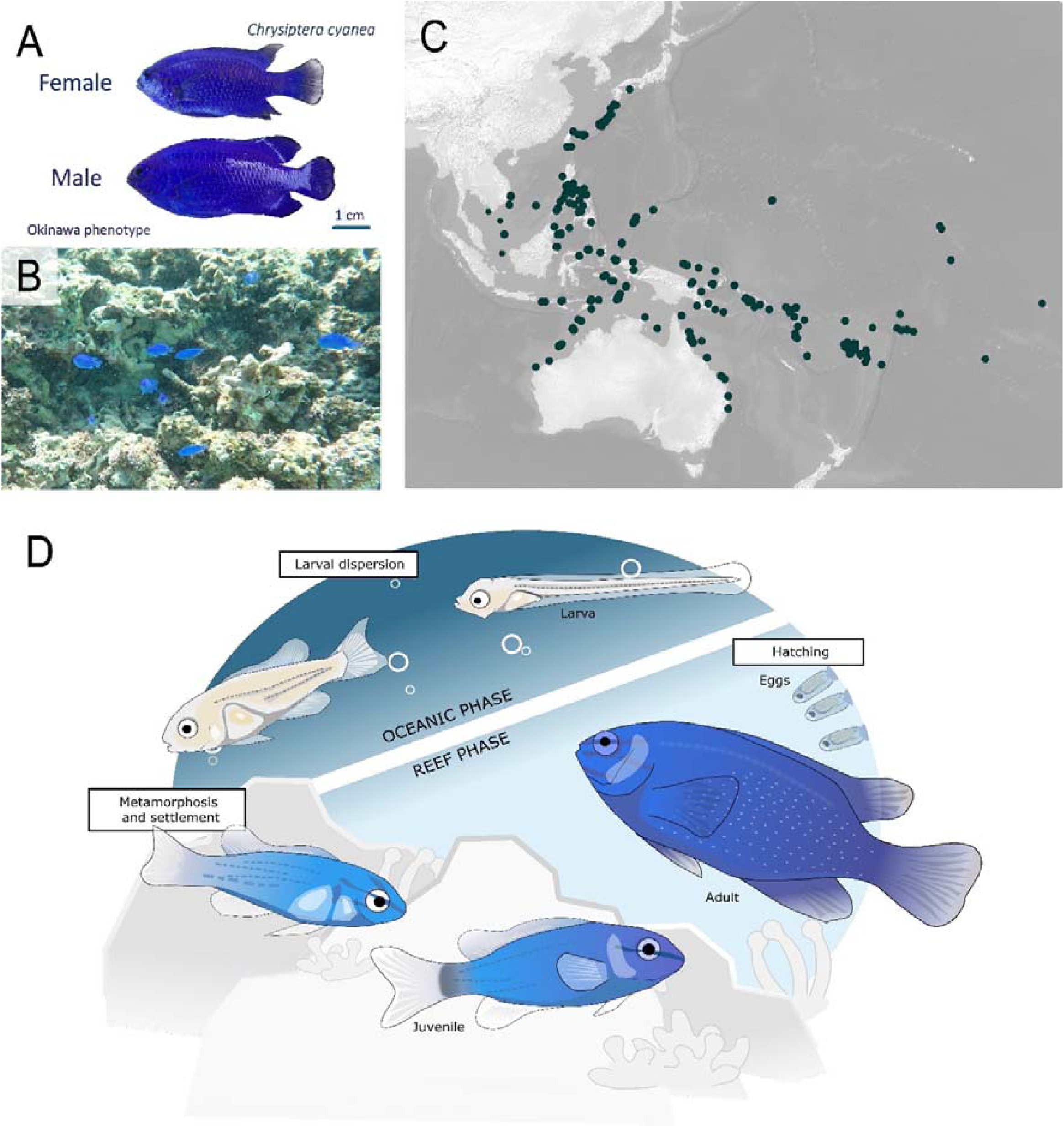
A. Female and male *Chrysiptera cyanea* collected in Sesoko beach, Okinawa, in September 2022. B. Underwater photography of a school of *C. cyanea* at Sesoko Beach, September 2022. C. Global georeferenced records of *C. cyanea* (477 records) extracted from the Wallace module [62] and plotted with QGIS 3.22.9 [63]. D. Life cycle of *C. cyanea* from hatching to the adult stage. Drawing by Stefano Vianello.

The life cycle of *C. cyanea* is akin to the majority of damselfish species, with a pelagic larval duration of 17-21 days [17] followed by a return to the reef coupled with the metamorphosis from larvae to juveniles. The juveniles then grow and eventually become a sexually mature female (Figure 1D). The presence of a sex-specific size distribution and bias in the sex ratio of *C. cyanea* on the reef is indicative of protogynous hermaphroditism, with females turning into males [14,18].

In Okinawa, females have transparent caudal fins and males have opaque blue caudal fins (Figure 1A). This phenotype is different from that of other locations in the Indo-Pacific, where males exhibit a yellow to orange tail. Studies of COI showed divergence between *C. cyanea* from Indonesian/Australian, Philippines, and Tonga which suggest the presence of allopatric divergence with possible new damselfish species (different COI reported in Tonga [19]. *C. cyanea* was originally described in Timor as a blue fish with yellow fins (Quoy and Gaimard 1825). A synonymous species, *Chrysiptera punctatoperculare* (Fowler, 1946) was described in Aguni, Okinawa. Here, we identified the species living in Okinawa, with blue fins, as *Chrysiptera cyanea* on the basis of a recent taxonomy guide [20].

This damselfish species is widespread along the coastal zones of Okinawa, making it an easily accessible study model for various behavioural, developmental, ecological, ecotoxicological questions. To tackle functional and molecular-level questions, in particular those based on gene expression, the availability of a genome is a considerable resource. For instance, in the case of transcriptomic studies, genome availability allows to perform reference-based alignments: such alignments are a robust method to examine RNA sequencing results while minimising issues such as incorrect assemblies of paralogous sequences (from gene duplication and diversification, leading to families of genes with similar sequences), preventing an underestimation of gene expression levels, as well as avoiding incomplete and fragmented assemblies which can be produced by *de novo* assembly methods [21]. To date, no genomic data is available for any *Chrysiptera* species. Here, we present a highly complete genome obtained from PacBio long-read sequencing, making this study the first to publish a genome for this coral reef fish genera, and adding to the number of genomes available for damselfishes. These genomes can also be useful to improve the analysis of the unique adaptations present in anemonefishes [22].

## Methods

### Fish collection and DNA sequencing

A male *Chrysiptera cyanea* (total length: 5.6cm) was collected using SCUBA and hand nets on Sesoko beach in Okinawa, Japan (26.6509 N, 127.8564 E) on September 30^th^, 2022. The fish was kept under natural conditions in a 270 L flow-through outdoor tank at the OIST Marine Science Station until October 19^th^, 2022. It was euthanised in a 200mg/L Tricaine Methanesulfonate (MS222) solution following the guidelines for animal use issued by the Animal Resources Section of OIST Graduate University. Liver tissue destined for genome sequencing was snap frozen in liquid nitrogen and stored at –80°C. Eye, gill, liver and muscle tissue from the same male fish were preserved in RNAlater for transcriptomic sequencing and stored at –30°C.

Genomic DNA was extracted from the snap-frozen liver using the Monarch HMW DNA Extraction Kit for tissues (New England BioLabs). Library preparation and sequencing were performed by the OIST Sequencing Section. First, the genomic DNA was sheared to 25kb using the Megaruptor 3 (Diagenode), as longer reads improve the ease of genome assembly. Library preparation was conducted using the SMRTbell Express Template Prep Kit 3.0 (PacBio, #102-182-700), following the manufacturer’s protocol. Library size selection was carried out using the BluePippin system (SageScience). After preparation with the Sequel Binding Kit 3.2 to bind the polymerase and sequencing primers, sequencing was performed on two SMRT cells with Circular Consensus Sequencing (CCS) runs using PacBio Sequel IIe HiFi (Pacific Biosciences, CA, USA).

RNA was extracted from the eye, gill, liver, and muscle tissue using the Maxwell® RSC simplyRNA tissue kits and an automated Maxwell® RSC instrument, following manufacturer recommendations (Promega, Cat. No. AS2000, Wisconsin, USA). The quality and concentration of RNA were assessed with an Invitrogen Qubit Flex benchtop fluorometer and an Agilent 4200 TapeStation. Library preparation was performed by the OIST Sequencing section using the NEBNext Ultra II Directional RNA library prep kit for Illumina (New England BioLabs, USA). Sequencing was performed using an Illumina NovaSeq6000 platform at OIST.

#### Sequencing data processing and genome assembly

The complete list of software, version, and parameters is provided in Table 1. Quality control was performed with FastQC 0.11.9 on the raw reads from each SMRT cell. The genome size was estimated based on a k-mer approach using Jellyfish 2.2.7 and GenomeScope 2.0. The DNA sequences were assembled with the Improved Phased Assembler 1.3.1 (IPA) [23] without phasing – *i.e.,* without separating the parental alleles into haplotypes, as this option yielded the best assembly results out of different IPA and Flye 2.9.1 [24] assemblies based on Quast 5.2.0 statistics [25], BUSCO (Benchmarking Universal Single-Copy Orthologs) scores (actinopterygii_odb10, BUSCO 4.1.2) [26], and similarity to the k-mer estimation of the genome size. Merqury 1.3 was used to estimate the genome completeness and error rate. Purge Haplotigs 1.1.3 was used to improve contiguity by removing allelic contigs from the non-phased IPA assembly [27]. The repeats were annotated with RepeatModeler 2.0.3 with the parameter –LTRStruct [28]. RepeatMasker 4.1.1 [29] was run to identify repetitive elements in the RepeatModeler output and the vertebrate library of Dfam. Repetitive elements were then softmasked with BEDTools 2.30.0 [30].

**Table 1.**
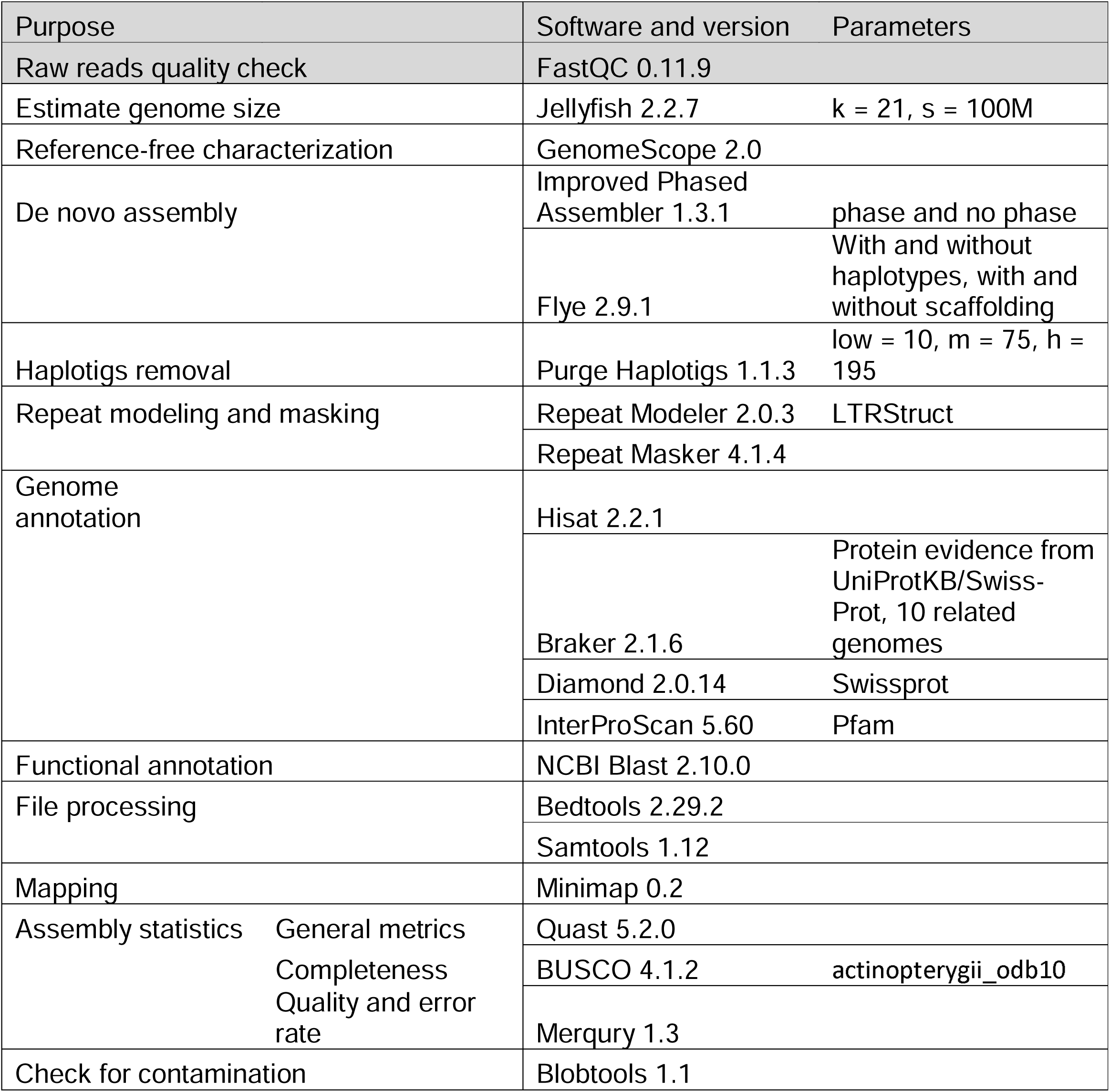
Software and parameters used for the genome assembly and annotation of *Chrysiptera cyanea*.

### Contig scaffolding on clownfish reference genomes

The chromosome-scale genome assemblies for *Amphiprion clarkii, A. ocellaris, A. percula, Acanthochromis polyacanthus,* and *Dascyllus trimaculatus* were used to map the 91 contigs of the *C. cyanea* assembly to the chromosomes using MUMmer 3.23 [31]. Dot plots were generated using ggplot2 [32] following data filtering to remove any alignments shorter than 10,000 bases [33].

### Transcriptome sequencing data processing

Quality control of the RNA sequencing data from the eye, gill, liver, and muscle tissue was performed with FastQC 0.11.9 [34]. Following this quality check, the sequences were processed by trimming the adapters used by Illumina on each transcript as well as dropping low quality sections with Trimmomatic 0.39 [35]. The transcriptomic reads were mapped to the contig sequences with HISAT2 2.2.1 [36], then converted to BAM format from SAM format with SAMtools 1.12 [37]. Lastly, the number of transcripts per gene was quantified using Kallisto 0.46.2 [38].

### Prediction of gene models

The position of protein-coding gene structures in the softmasked assembly was predicted by BRAKER 2.1.6 [37,39–50] using the transcriptome of the eye, gill, liver, and muscle of the fish used for the genome. The position of the protein-coding gene structures was also predicted using protein evidence from UniProtKB/Swiss-Prot [51] as well as selected fish proteomes from the NCBI database (*Amphiprion ocellaris*: 48,668 sequences, *A. clarkii*: 25,025, *Danio rerio*: 88,631, *Acanthochromis polyacanthus*: 36,648, *Oreochromis niloticus*: 63,760, *Oryzias latipes*: 47,623, *Poecilia reticulata*: 45,692, *Stegastes partitus*: 31,760, *Takifugu rubripes*: 49,529, and *Salmo salar*: 112,302). The genes with evidence from the transcriptome or protein hints, and with homology to the Swiss-Prot protein database and Pfam domains identified with Diamond [42] and InterProScan [52], were selected. NCBI BLAST 2.10.0 [53] was used to perform the functional annotation of the final gene models.

The output was assessed using Quast 5.2.0 [25] as well as by calculating the number of BUSCO genes present in the assembly [26].

### Gene expression analysis

Using the R package tispec, the tissue-specify index (τ) was calculated for each gene using the R package tispec v0.99 [54]. The relationship between τ and the transcript per million (TPM) gene expression values visualized on a 2D histogram with ggplot2 [32]. An upset plot from UpSetR v1.4.0 [55] was used to visualise the TPM values per tissue (brain, eye, gill, liver, muscle).

## Results

### Genome assembly of C. cyanea

We assembled the genome of the damselfish *C. cyanea* by sampling one individual from Okinawa and obtaining 3,335,935 PacBio reads. The average read length was 25,387 for a total number of 84,688,690,513 sequenced bases. FastQC 0.11.9 did not detect low quality reads (Supplementary Figure 1).

Multiple *de novo* assembly options were tested (Table 2) using Flye and the Improved Phased Assembler (IPA). The IPA no-phasing primary assembly was chosen as it yielded the highest N50 (30.6 Mb), lowest number of contigs (103), and a genome length of 899.7 Mb. From the raw reads, the genome size had been estimated with a k-mer approach to be 753-754Mb (using Jellyfish and GenomeScope with k-mer sizes 21 and 31; Table S1, Figure S4). This is 16% lower than the assembly; the discrepancy might be due to the high repeat content of teleost fish genomes, leading to an underestimation of the genome size using a k-mer approach. Similarly, the estimated size of the genome using the k-mer approach was lower than the final assembly for *A. ocellaris* [4]*, A. clarkii* [9] and *D. trimaculatus* [3]. The assembly size was close to the reported lengths of other damselfish genomes. For instance, the genome size of *Amphiprion ocellaris* is 861.4 Mb [4], *A. clarkii* is 843.6 Mb [9], *A. percula* is 908.8 Mb [1], and *D. trimaculatus* is 910.8 Mb [3].

**Table 2.**
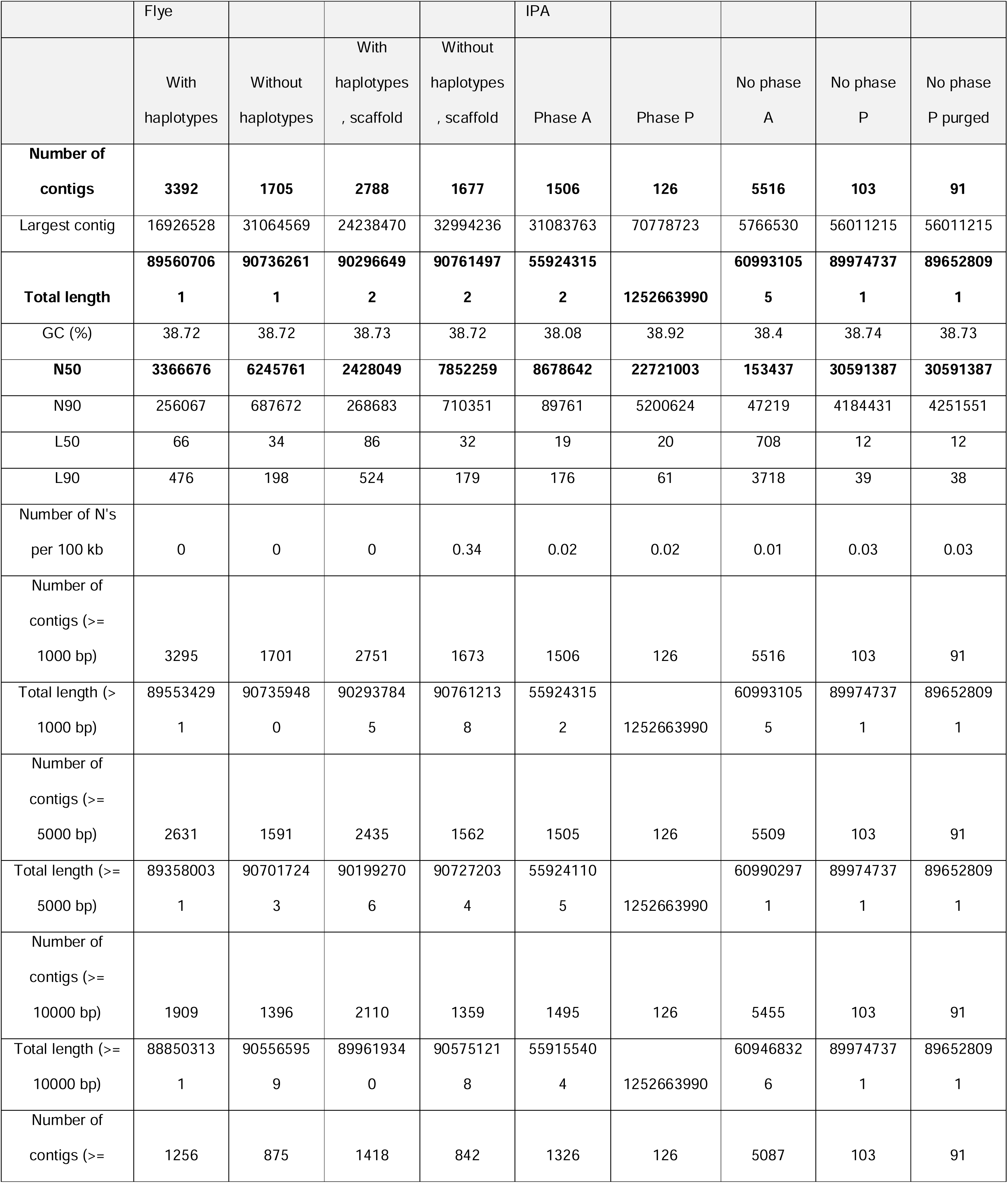
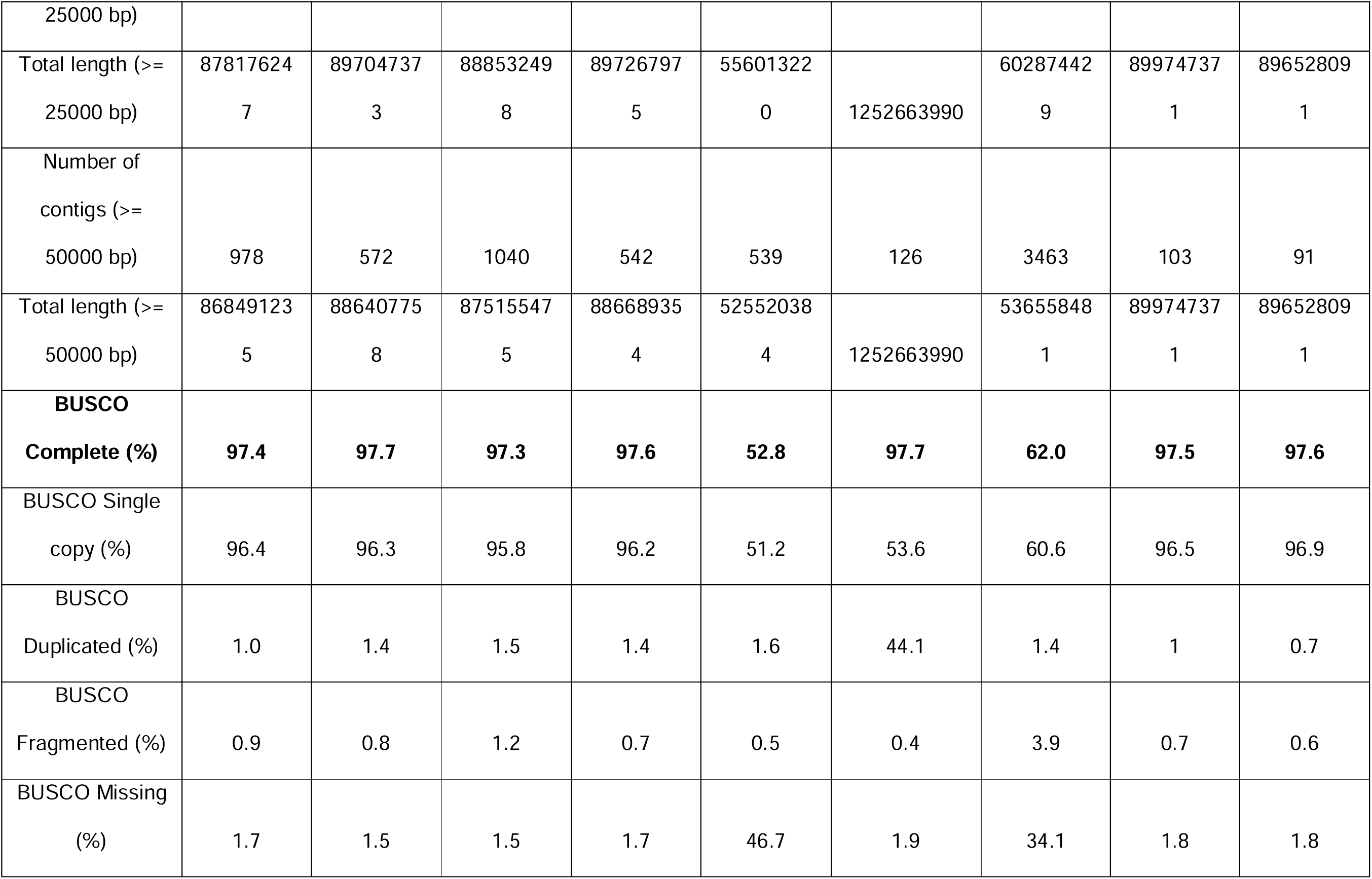
Statistics generated with Quast 5.2.0 and BUSCO 4.1.2 about the different primary genome assemblies produced by the Flye 2.9.1 and the Improved Phased Assembler 1.3.1 (IPA), as well as about the final assembly, which was obtained from the non-phased purged assembly (read depth cutoff: low = 10, medium = 70, high = 195) generated with Purge Haplotigs 1.1.3, with softmasked repeats obtained from Repeat Modeler 2.0.3 and Repeat Masker 4.1.4.

The *de novo* assembly was curated using Purge Haplotigs (removing one contig when syntenic pairs of contigs are detected) [27]. 12 contigs were removed to obtain a curated assembly with 91 contigs, of a total length of 896.5Mb. The GC content of the genome was 38.73% and the mean base-level coverage was 94.5x (Table 2). 97.6% of BUSCO genes were retrieved in the final genome assembly, with 96.9% single copy and 0.7% duplicated. 0.6% of BUSCO genes were found to be fragmented and 1.8% were missing. The completeness of the genome estimated by Merqury was 90.129% with a mean QV of 48.9 and error rate of 0.0000138 (one nucleotide error per 72.6kb).

324,222,639 bp (36.16% of sequences) were identified as being repeat content using RepeatMasker, with 20% DNA repeat elements, 18% simple repeats (microsatellites), 6% long interspersed nuclear elements, 2% low complexity repeats, 2% long terminal repeats and under 1% each of short interspersed nuclear elements, RC (rolling circle), tRNA, retroposon, rRNA, snRNA, and Satellite elements. 50% of the repetitive elements in the *C. cyanea* genome could not be identified.

As a reference, a comparison of key statistics about this genome assembly with previously published chromosome-scale genomes of Pomacentrid fish is provided in Table S3. Overall, while our coverage was slightly lower than other genomes, we were able to obtain a much more contiguous *de novo* assembly (91 contigs *vs.* 951 in *A. polyacanthus* and over 1400 in all other assemblies) by using very long 25kb DNA fragments for sequencing (rather than the recommended 10-20kb). Our contig N50 of 30.6Mb was six to thirty times larger than those of the other assemblies. The GC, repeat contents, and BUSCO completeness scores were similar to the other genomes.

The 91 contigs were mapped against the chromosome-scale genomes of *Amphiprion clarkii, A. ocellaris, A. percula, Dascyllus trimaculatus,* and *Acanthochromis polyacanthus* using MUMmer. After filtering to remove aligned segments shorter than 10,000 bases, over 40% of the bases of the assembly of *C. cyanea* were matched with the reference genomes of *Amphiprion clarkii, A. ocellaris, A. percula,* and *Acanthochromis polyacanthus*. However, only 20.3% were retrieved in the alignment with *Dascyllus trimaculatus* (Table S5); the latter genome was reported to have undergone rearrangements [3].

In total, 48 contigs were mapped onto the reference genomes (filtering out alignments shorter than 10,000 bases). The correspondences are provided in Table S4. 16, 18, 14, 16, and 0 contigs had over 50% of the bases aligning to the reference genomes of *Amphiprion clarkii, A. ocellaris, A. percula, Acanthochromis polyacanthus,* and *Dascyllus trimaculatus* (Figure 2A). Forty-six contigs were aligned with a single chromosome (Figure 2A). Forty-three contigs did not robustly map to any chromosome of the reference genomes, among which contig 36 was the largest contig that could not be matched (4.6Mb).

**Figure 2.**
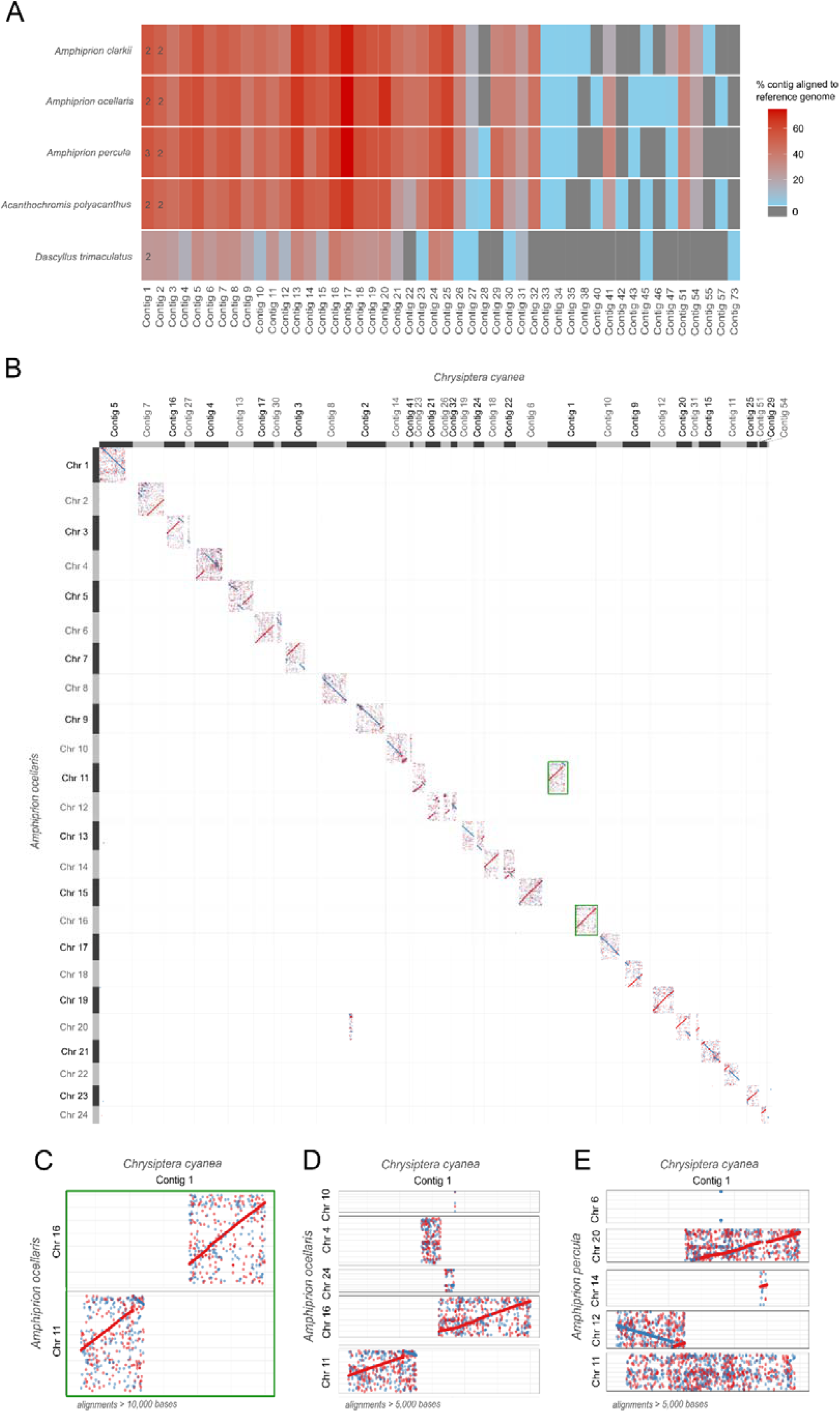
A. Percentage of bases in each contig aligning to the reference genomes. Only contigs aligning to at least one chromosome, with at least 10,000 bases retrieved from the reference genome, are displayed (48 of the total 91 contigs). Based on this threshold, most contigs align to only one reference chromosome, with exceptions indicated by a number inside of the corresponding tile. Contig 1 is indicated with a green frame. B. Dotplot of the alignment of the chromosome scale *Amphiprion ocellaris* genome with the contigs from *Chrysiptera cyanea* (alignments longer than 10,000 bases). This dotplot is a larger version of the green frame on panel A. C. Dotplot of the alignment of contig 1 from Chrysiptera cyanea with chromosome 11 and 16 of *A. ocellaris* (alignments longer than 10,000 bases). D. Dotplot of the alignment of contig 1 from Chrysiptera cyanea with chromosome 4, 10, 11, 16, and 24 of *A. ocellaris* (alignments longer than 5,000 bases). E. Dotplot of the alignment of contig 1 from *Chrysiptera cyanea* with chromosomes 6, 11, 12, 14, and 20 of *Amphiprion percula* (alignments longer than 5,000 bases).

**Figure 3.**
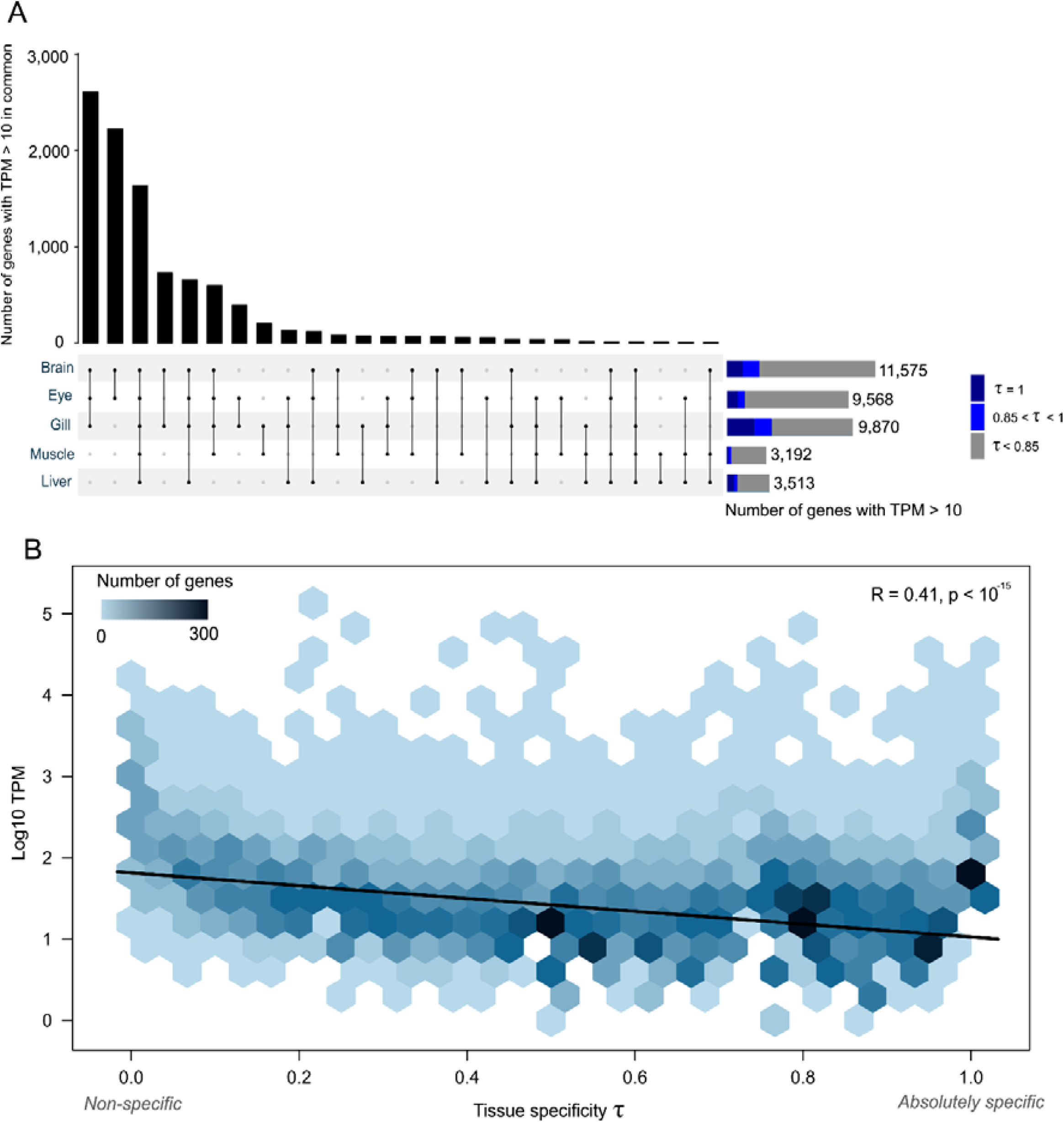
A. Upset plot displaying the number of genes expressed (intersection size) in individual and combinations of different tissues (brain, eye, gill, liver, muscle). Transcripts per million (TPM) values above 10 were used as a threshold for gene expression. B. Two-dimensional histogram displaying the relationship between the maximum TPM (log-transformed) and tissue specificity index (Tau, τ) of each gene. Trendline displays Pearson’s correlation between τ and log10.

Two *Chrysiptera cyanea* contigs were mapped to more than one chromosome from the reference assemblies: contigs 1 and 2 (the two largest contigs, of sizes 56.0Mb and 45.2Mb). This may indicate a chimeric assembly in the analysis of the data or genomic rearrangement. For contig 1, the alignment against the *A. ocellaris* genome highlighted a missing zone between the 19,943,083 and 33,394,537^th^ bases with no alignments longer than 10,000 bases (Figure 2C). Similar results were found for comparisons with the other reference genomes. Lowering the filtering threshold to include alignments between 5,000 and 10,000 bases against *A. ocellaris* retrieved some relatively scrambled alignments in the gap mentioned above (Figure 2D). This could indicate a chimeric assembly of contig 1, which may in fact consist of two chromosomes in *C. cyanea.* However, the comparison with *A. percula* for alignments longer than 5,000 bases found that the missing zone maps onto *A. percula*’s chromosome 20 (Figure 2E). This may point towards genomic rearrangement from two or more ancestral chromosomes, with divergence through evolution leading to the low mapping rate between 19Mb and 33Mb on contig 1.

### *C. cyanea* gene annotation

The genome was annotated using BRAKER v.2.1.6 based on mRNA and protein evidence, leading to the prediction of 46,873 gene models. These were filtered to only keep the longest isoform of each gene model. Only genes with mRNA evidence, or homology to the Swiss-Prot protein database and Pfam domains were kept. The completeness of the final set of 28,173 genes was assessed using BUSCO, with a final score of 96.6%, including 96.0% single copy genes and 0.6% duplicated genes. 1.5% BUSCO genes were fragmented, and 1.9% were missing from the final transcriptome. Of the final set of 28,173 genes, 19,356 (69%) had at least one associated GO term. Lastly, 1,802 genes (6.4% of all genes) were located on contigs that could not be matched to any chromosome from the five reference genomes.

### Tissue specificity of the gene expression

The tissue specificity of the 28,173 identified genes, or “transcriptomic atlas” of *Chrysiptera cyanea,* was assessed using the transcriptomes of the eye, gill, liver, and muscle of the fish used for the genome assembly, as well as the brain from another fish collected on the same day and site. In total, 1,639 genes were expressed in all five tissues (with transcripts per million (TPM) values above 10 in each tissue), which was similar to previous reports in anemonefish [4,56]. The tissue specificity index τ was calculated for each gene (for which τ near 0 indicates broadly expressed genes, and τ near 1 specific genes) [57,58]. 3,090 genes had a τ value below 0.2, which corresponds to housekeeping genes (similar expression levels across most tissues without bias) [57]. This number was relatively similar to those reported for *Amphiprion* species, notably for *A. clarkii* (3,697 genes) [56] and *A. ocellaris* (3,431 genes) [4]. Following log-transformation and normalisation, 7,396 genes with τ values beyond 0.85, indicative of high tissue specificity, were identified. Of these, 4,258 genes were absolutely specific with a τ = 1 (expressed in only one type of tissue). This is a relatively high number and may be linked to the low amount of tissue types considered here, compared to other studies of damselfish (five here against *e.g.,* twelve in [56]). The gill tissue had the highest number of highly specific genes, with 1,170 genes with τ = 1, followed by the brain tissue (1,094), eye (780), liver (480), and muscle (134). These values were proportional to the total number of genes expressed in each tissue, which was higher in the brain, eye, and gill and lower in the liver and muscle, similarly to other damselfish. Lastly, tissue-specific genes tended to have lower expression levels (negative correlation of τ value with gene expression level, R = 0.41, p < 10^-15^), as previously reported for *A. clarkii* [56] and *A. ocellaris* [4].

## Discussion

The Sapphire Devil *Chrysiptera cyanea* is a widely distributed damselfish in the Indo-Pacific area. In Okinawa, it is among the most common species on reef flats [59]. It has been mainly studied to elucidate the roles of various environmental controls on their reproduction, and investigate related hormonal processes [12,15,16]. To further the potential of biomolecular analyses based on this species, this study generated the first genome of a *Chrysiptera* fish, from a male individual collected in Okinawa, Japan. This genome will be of high value for future genetic-based approaches, from population structure to gene expression analyses. Among hot topics in research, the difference between anemonefish and other damselfish is particularly examined. Here, we provide a new high quality non-anemonefish genome which will be of high relevance to further the depth of such analyses.

Using PacBio HiFi, we assembled a high-quality genome of size 896.5Mb, within the range of other related species, with a high completeness of 97.6% BUSCO genes. Of particular interest, although Hi-C was not generated for this study, the use of PacBio HiFi long-read technology, the Improved Phased Assembler, and Purge Haplotigs allowed us to identify 91 contigs. This is a low number of contigs, particularly compared to those for other reef fish with scaffold-level genomes based on Illumina short read approaches. The *Chrysiptera cyanea* genome contigs closely mapped against chromosome-scale genomes from other Pomacentrids, with several homologous contig-chromosome pairs (Figure 2, Table S6). The high quality of the genome assembled here is indicative that the sequencing depth, choice of technology, and downstream pipelines were sufficient to retrieve long contigs and achieve a near chromosome-scale genome assembly.

Several architectural variations in the genome of *Chrysiptera cyanea* were detected by comparison with the five reference genomes. Some contigs showed homology with their reference chromosomes (*e.g*., > 10,000-base segments from contig 8, representing 55.8% of the contig length, were homologous with over 50% of the bases of chromosome 8 from *A. percula*, Figure 2B, Table S5, S6). However, others showed rearrangements that ranged from inversions with or without frame shifts to a possible rearrangement of ancestral chromosomes in the Cheiloprionini lineage (Figure 2B). In particular, over 50% the bases of contig 1 were aligned to at least two chromosomes of all reference genomes, with about half of the bases mapping to one chromosome and half to another (Figure 2, Table S7). No *Chrysiptera cyanea* contigs robustly scaffolded to chromosome 24 of *Acanthochromis polyacanthus* and *Dascyllus trimaculatus* (Table S4). As chromosome 24 of *D. trimaculatus* was reported to be highly rearranged [3], the absence of alignment with this chromosome highlights that the rearrangement occurred in the Chrominae branch, which separated from the Pomacentrinae branch, to which *Amphiprion, Acanthochromis,* and *Chrysiptera* belong, over 50 million years ago [60,61]. The relative distance of *Chrysiptera* to *Dascyllus trimaculatus* compared with the other reference species is also illustrated by contig 18, which is the contig showing the highest rate of alignment with most genomes. Indeed, 67.7 to 75.1% of the bases of contig 18 were aligned to the *Amphiprion clarkii, A. ocellaris, A. percula* and *Acanthochromis polyacanthus* reference genomes (e.g., Figure 2F with *A. percula*) but only 34% with *Dascyllus trimaculatus* (Figure 2G). Lastly, the similarity in results when comparing the *Chrysiptera cyanea* assembly with each of the four *Amphiprion* and *Acanthochromis* reference genomes can also be linked to the evolutive history of Pomacentrids. *Chrysiptera cyanea* is part of the Cheiloprionini, which split from the other Pomacentrinae, notably from the branch to which *Amphiprion* and *Acanthochromis* belong, over 35 million years ago, leading to similar differences in the comparisons to all of these genomes [60,61].

## Conclusion

In this study, the first genome assembly for a *Chrysiptera* species was generated with 91 contigs. The contigs were successfully aligned to related reference genomes, allowing for forays into the genomic architecture of Pomacentrids with respect to their evolutive relationship. *Chrysiptera cyanea* is easy to breed and maintain in aquaria and is highly abundant in Indo-Pacific coastal waters, particularly in Okinawa, Japan. The generation of this high-quality genome will further the potential of this species as a coral reef model species for research questions requiring biomolecular approaches.

## Data availability

The genome and transcriptome sequencing reads are deposited in the NCBI GenBank database under the BioProject PRJNA1167451. The genome assembly, annotation, and proteome for *C. cyanea,* as well as BUSCO outputs and detailed scripts for each genome assembly and annotation step, are available on FigShare: https://figshare.com/s/6b6be390dc8b4c5d025b.

## Acknowledgements

The authors would like to thank the OIST Sequencing Section for the library preparation and sequencing, as well as Stefano Vianello for providing the drawings of *Chrysiptera cyanea*.

## Declarations

The experimental procedure was conducted in accordance with ethical procedures recommended by the Okinawa Institute of Science and Technology Graduate University. The authors declare no competing interests.

## Supplementary Materials

**Table S1.** Summary table of the estimation of the genome size based on k-mers using Jellyfish 2.2.7 and GenomeScope 2.0.

|  | k-mer size = 21 |  | k-mer size = 31 |  |
| --- | --- | --- | --- | --- |
|  | min | max | min | max |
| Heterozygosity | 0.797429% | 0.804408% | 0.540891% | 0.545631% |
| Haploid length | 753,509,729 | 754,193,212 | 753,509,728 | 754,193,211 |
| Repeat length | 84,198,896 | 84,275,270 | 84,198,892 | 84,275,266 |
| Unique length | 669,310,832 | 669,917,941 | 669,310,836 | 669,917,945 |
| Model fit | 90.2553% | 91.4736% | 90.2553% | 91.4736% |
| Read error rate | 0.267665% | 0.267665% | 0.181441% | 0.181441% |

**Table S2.** Number of nucleotide bases and genes on each contig of the *Chrysiptera cyanea* genome assembly.

| Contig name | Size | Number of genes |
| --- | --- | --- |
| 1 | 56011215 | 1691 |
| 2 | 45244965 | 1482 |
| 3 | 41200874 | 1262 |
| 4 | 39499864 | 1440 |
| 5 | 38661106 | 1015 |
| 6 | 37342079 | 1124 |
| 7 | 36429693 | 1157 |
| 8 | 35288363 | 1278 |
| 9 | 31922934 | 1069 |
| 10 | 30696959 | 926 |
| 11 | 30671322 | 852 |
| 12 | 30591387 | 1229 |
| 13 | 29350308 | 1029 |
| 14 | 28011985 | 1059 |
| 15 | 25310964 | 677 |
| 16 | 24774977 | 902 |
| 17 | 23869012 | 892 |
| 18 | 22772264 | 789 |
| 19 | 19058377 | 512 |
| 20 | 18402649 | 769 |
| 21 | 17741195 | 503 |
| 22 | 14111127 | 360 |
| 23 | 14090289 | 329 |
| 24 | 12221435 | 427 |
| 25 | 11992873 | 425 |
| 26 | 11817260 | 301 |
| 27 | 10508735 | 311 |
| 28 | 9326607 | 270 |
| 29 | 8821376 | 182 |
| 30 | 8416732 | 227 |
| 31 | 7840817 | 236 |
| 32 | 7505997 | 194 |
| 33 | 5628431 | 77 |
| 34 | 5502750 | 145 |
| 35 | 4884161 | 154 |
| 36 | 4641543 | 131 |
| 37 | 4409973 | 96 |
| 38 | 4251551 | 114 |
| 39 | 4184431 | 71 |
| 40 | 4017153 | 131 |
| 41 | 3905345 | 105 |
| 42 | 3763230 | 135 |
| 43 | 3622033 | 113 |
| 44 | 3616284 | 73 |
| 45 | 3507541 | 69 |
| 46 | 3369239 | 85 |
| 47 | 2871160 | 85 |
| 48 | 2816112 | 38 |
| 49 | 2768566 | 91 |
| 50 | 2746381 | 136 |
| 51 | 2497950 | 94 |
| 52 | 2456027 | 67 |
| 53 | 2331100 | 36 |
| 54 | 2034662 | 42 |
| 55 | 1988681 | 65 |
| 56 | 1890354 | 36 |
| 57 | 1851746 | 30 |
| 58 | 1825140 | 44 |
| 59 | 1819243 | 79 |
| 60 | 1756753 | 54 |
| 61 | 1755205 | 75 |
| 62 | 1735356 | 16 |
| 63 | 1580605 | 76 |
| 64 | 1562090 | 72 |
| 65 | 1508740 | 28 |
| 66 | 1401674 | 60 |
| 67 | 1392888 | 20 |
| 68 | 1369576 | 19 |
| 69 | 1360515 | 61 |
| 70 | 1223198 | 26 |
| 71 | 1205908 | 19 |
| 72 | 1096930 | 79 |
| 73 | 1046040 | 8 |
| 74 | 993160 | 41 |
| 75 | 946463 | 58 |
| 76 | 831928 | 20 |
| 77 | 690090 | 11 |
| 78 | 662047 | 32 |
| 79 | 534994 | 23 |
| 80 | 507718 | 34 |
| 81 | 497595 | 11 |
| 82 | 474921 | 8 |
| 83 | 464461 | 14 |
| 84 | 344082 | 14 |
| 85 | 205924 | 6 |
| 86 | 185716 | 5 |
| 87 | 182094 | 11 |
| 88 | 98395 | 5 |
| 89 | 77521 | 2 |
| 90 | 73239 | 1 |
| 91 | 55738 | 3 |

**Table S3.**
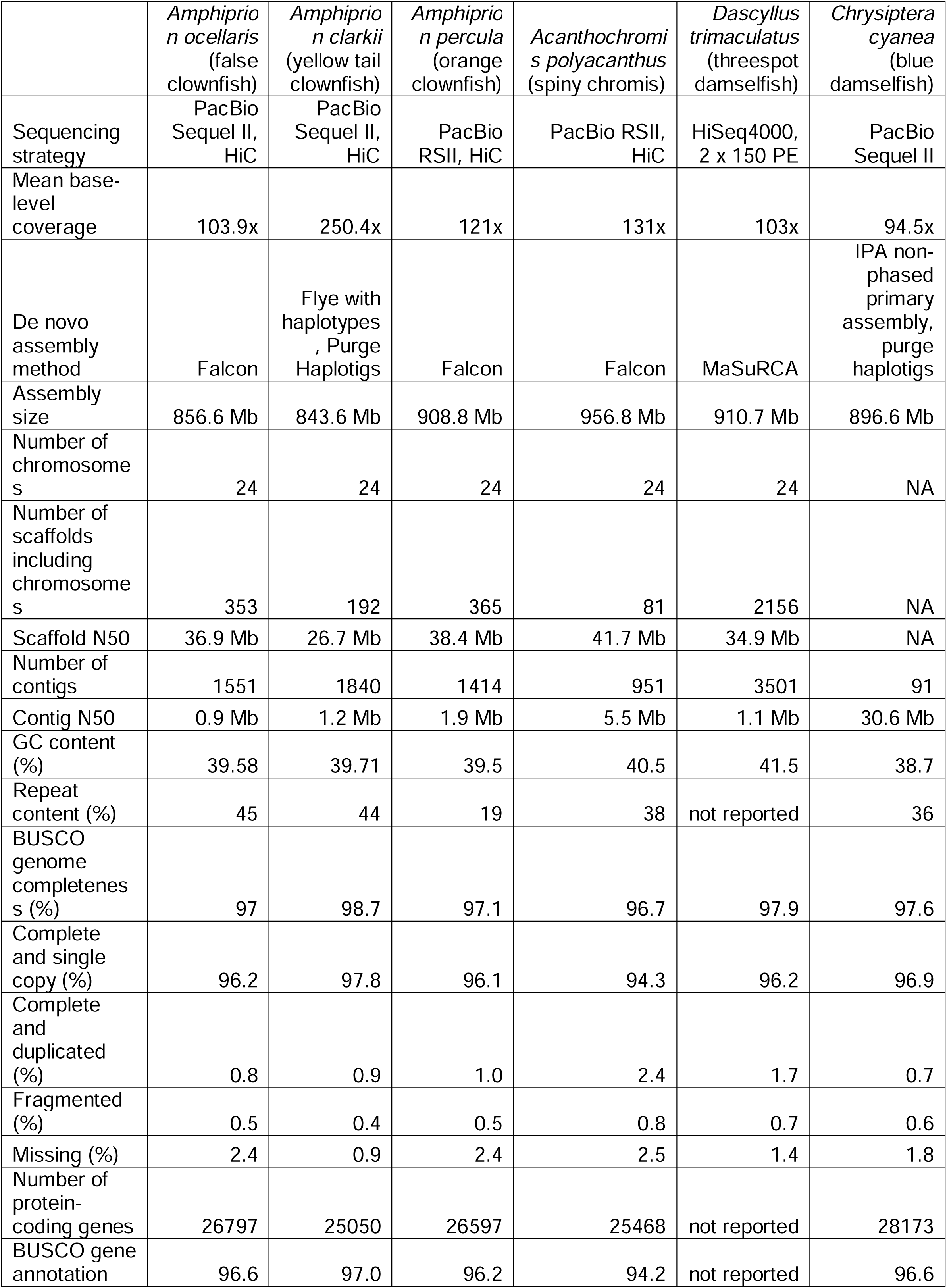
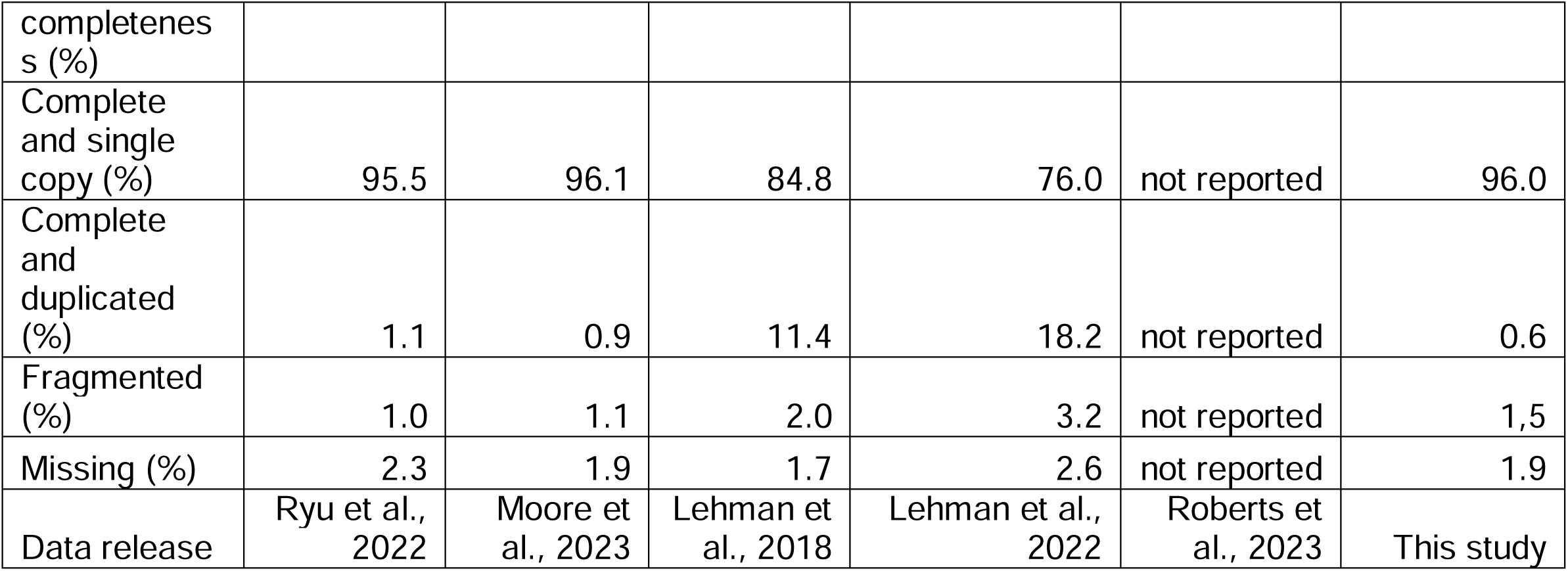
Comparison of the genome assembly strategy and output for *C. cyanea* with other chromosome-scale genome assemblies of Pomacentrid fish.

**Table S3.**
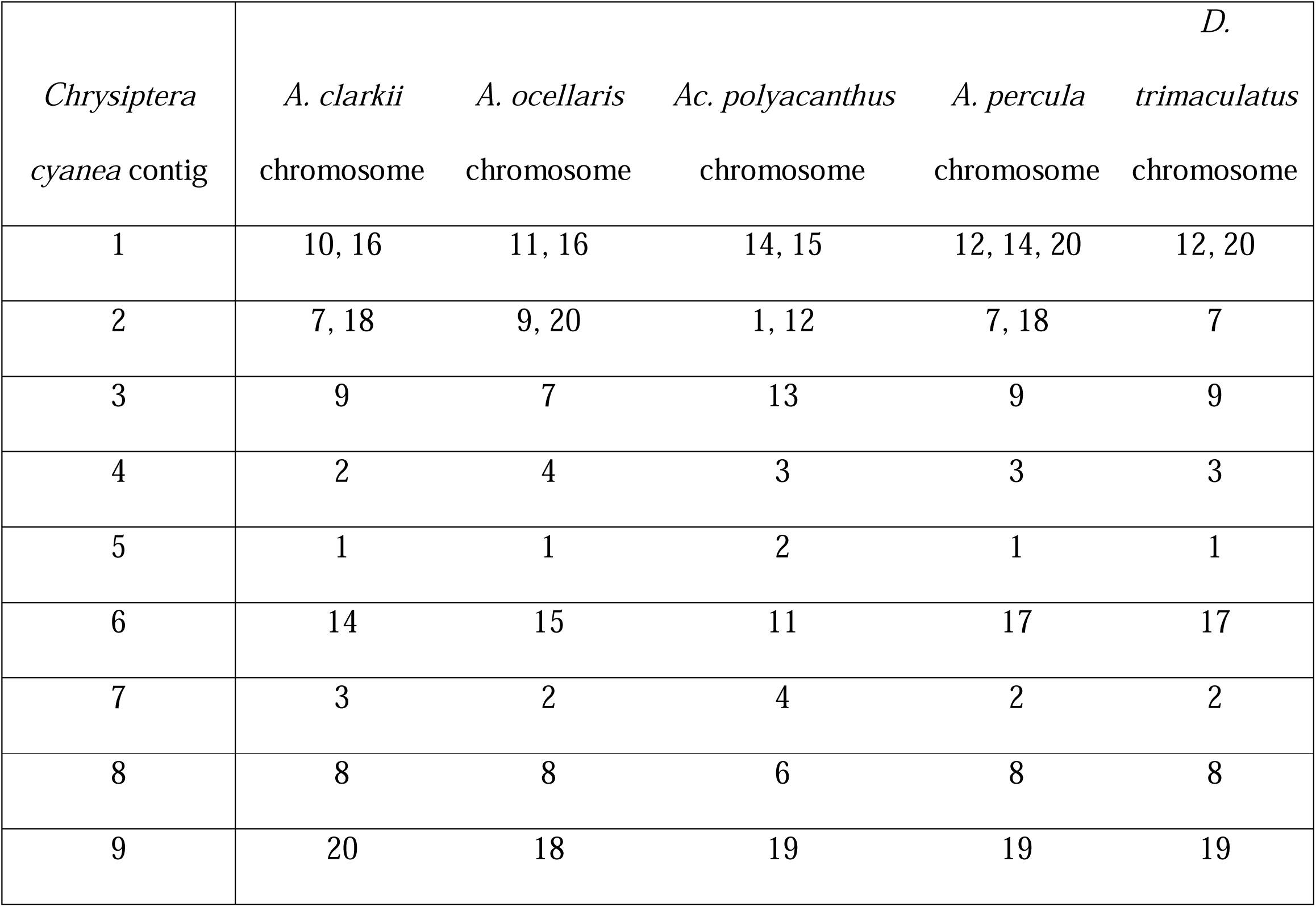
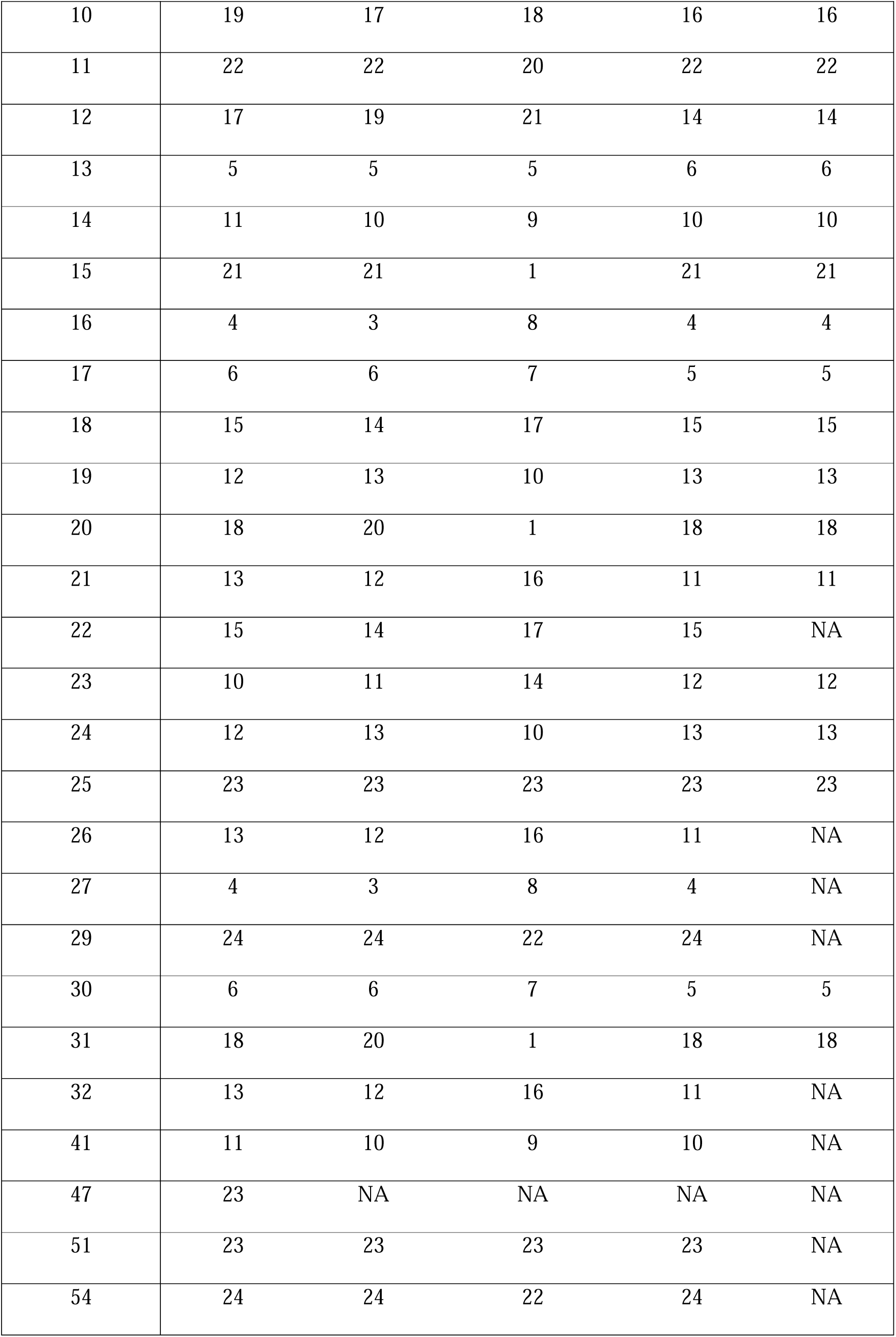
Correspondence of the contigs of the genome assembly for *C. cyanea* and the chromosomes of the reference genomes of *Dascyllus trimaculatus, Acanthochromis polyacanthus, Amphiprion clarkii, Amphiprion ocellaris,* and *Amphiprion percula*. 223 chromosome-contig matches were retrieved. Matches with less than 100,000 bases are not indicated in the table (50 matches removed). Scaffolding was performed using the nucmer function of the MUMmer 3.23 package [31] removing alignments shorter than 10,000 bases.

**Table S4.** Percentage of *Amphiprion clarkii, A. ocellaris, A. percula, Acanthochromis polyacanthus,* and *Dascyllus trimaculatus* chromosome-scale genomes aligned with *Chrysiptera cyanea* contigs based on MUMmer 3.23. Alignments shorter than 10,000 bases were dropped.

| Chromosome | A. clarkii | A. ocellaris | A. percula | A. polyacanthus | D. trimaculatus |
| --- | --- | --- | --- | --- | --- |
| 1 | 52.9 | 51.8 | 49.2 | 37.5 | 29.5 |
| 2 | 47.2 | 47.6 | 45.5 | 44.6 | 30.7 |
| 3 | 47.8 | 40.8 | 51.1 | 42.0 | 16.3 |
| 4 | 44.2 | 54.2 | 41.9 | 37.5 | 26.8 |
| 5 | 47.8 | 48.2 | 52.0 | 39.2 | 21.5 |
| 6 | 52.1 | 53.7 | 45.0 | 42.8 | 27.4 |
| 7 | 56.7 | 45.0 | 54.4 | 41.6 | 32.9 |
| 8 | 46.8 | 51.8 | 50.7 | 36.2 | 29.6 |
| 9 | 46.3 | 56.1 | 49.6 | 37.6 | 26.6 |
| 10 | 56.1 | 46.1 | 35.4 | 40.1 | 21.9 |
| 11 | 46.7 | 55.1 | 43.4 | 40.0 | 9.1 |
| 12 | 48.7 | 45.0 | 53.2 | 49.4 | 20.0 |
| 13 | 44.2 | 47.9 | 46.6 | 45.2 | 27.1 |
| 14 | 50.9 | 53.9 | 48.3 | 47.5 | 12.6 |
| 15 | 54.7 | 51.3 | 53.1 | 40.8 | 19.5 |
| 16 | 49.5 | 51.1 | 41.9 | 29.3 | 10.0 |
| 17 | 37.0 | 45.4 | 49.7 | 39.0 | 29.5 |
| 18 | 42.7 | 39.3 | 39.9 | 41.0 | 21.1 |
| 19 | 44.8 | 48.3 | 35.4 | 39.8 | 22.7 |
| 20 | 45.1 | 49.4 | 46.4 | 37.8 | 23.0 |
| 21 | 46.9 | 46.9 | 44.6 | 44.3 | 13.1 |
| 22 | 47.9 | 46.3 | 46.0 | 14.9 | 29.2 |
| 23 | 34.7 | 32.4 | 31.1 | 27.9 | 16.0 |
| 24 | 17.8 | 17.5 | 19.5 |  |  |
| Total | 48.2 | 47.6 | 45.5 | 39.4 | 22.8 |

**Table S5.** Percentage of the *Chrysiptera cyanea* contigs aligned with *Amphiprion clarkii, A. ocellaris, A. percula, Acanthochromis polyacanthus,* and *Dascyllus trimaculatus* chromosome-scale genomes based on MUMmer 3.23. Alignments shorter than 10,000 bases were dropped.

| Contig | A. clarkii | A. ocellaris | A. percula | A.<br>polyacanthus | D. trimaculatus |
| --- | --- | --- | --- | --- | --- |
| 1 | 55.7 | 56.7 | 56.4 | 53.6 | 26.3 |
| 2 | 51.3 | 50.6 | 51.7 | 48.9 | 27.1 |
| 3 | 42.8 | 42.6 | 47.2 | 45.5 | 24.5 |
| 4 | 50.4 | 56.2 | 56.1 | 50.1 | 16 |
| 5 | 58.2 | 58.9 | 58.7 | 56.2 | 31.6 |
| 6 | 48.8 | 48.5 | 47.2 | 44.7 | 26.5 |
| 7 | 53.6 | 54.6 | 54.3 | 47.6 | 31.7 |
| 8 | 50.4 | 55.8 | 56.6 | 54.2 | 30.8 |
| 9 | 44.7 | 41.3 | 38 | 41.3 | 21.3 |
| 10 | 48.1 | 50.1 | 49.9 | 48.3 | 10.4 |
| 11 | 44.2 | 43.6 | 44.7 | 40.7 | 28.5 |
| 12 | 40 | 52.8 | 52.7 | 47.1 | 12.8 |
| 13 | 65 | 66.1 | 64.8 | 60.3 | 35 |
| 14 | 58.7 | 57.3 | 43.7 | 52.6 | 28.1 |
| 15 | 53.2 | 54.2 | 54.2 | 51.4 | 15.1 |
| 16 | 64.8 | 61.6 | 65.1 | 62.2 | 40.6 |
| 17 | 72.4 | 74.9 | 75.1 | 67.7 | 34 |
| 18 | 59.7 | 57.2 | 60.3 | 54.3 | 29.8 |
| 19 | 55.8 | 56.3 | 56.3 | 53.6 | 28.7 |
| 20 | 56.5 | 68.9 | 57.2 | 54.9 | 32.1 |
| 21 | 46 | 49.7 | 49.6 | 27.9 | 17.7 |
| 22 | 41.2 | 44.4 | 43.7 | 20.3 | NA |
| 23 | 45.3 | 44.5 | 45.3 | 38.2 | 1.8 |
| 24 | 56.2 | 56 | 55.8 | 53.6 | 37.9 |
| 25 | 61.3 | 63.2 | 63.1 | 56.3 | 34.6 |
| 26 | 36.6 | 37.2 | 39.3 | 27.3 | 0.1 |
| 27 | 15.6 | 15.1 | 14.7 | 2.1 | 0.1 |
| 28 | NA | NA | 0.2 | 0.1 | NA |
| 29 | 35.6 | 38.3 | 47.3 | 47.5 | NA |
| 30 | 38.1 | 38.2 | 38.3 | 23.9 | 1.4 |
| 31 | 26.3 | 26.5 | 26.3 | 17.3 | 13 |
| 32 | 45.3 | 44.2 | 43.1 | 44 | NA |
| 33 | 0.2 | 0.3 | 0.2 | 0.2 | NA |
| 34 | 0.3 | 0.3 | 0.2 | 0.3 | NA |
| 35 | 0.4 | 0.4 | 0.4 | NA | NA |
| 36 | NA | NA | NA | NA | NA |
| 37 | NA | NA | NA | NA | NA |
| 38 | 0.4 | NA | NA | NA | NA |
| 39 | NA | NA | NA | NA | NA |
| 40 | NA | 0.3 | NA | 0.3 | NA |
| 41 | 17.5 | 30.2 | 36.1 | 25.6 | NA |
| 42 | NA | NA | NA | 0.3 | NA |
| 43 | NA | 0.7 | 0.3 | NA | NA |
| 44 | NA | NA | NA | NA | NA |
| 45 | 1.6 | 0.3 | NA | 0.4 | 0.3 |
| 46 | NA | 0.4 | NA | NA | NA |
| 47 | 22 | 1 | 0.7 | 0.8 | NA |
| 48 | NA | NA | NA | NA | NA |
| 49 | NA | NA | NA | NA | NA |
| 50 | NA | NA | NA | NA | NA |
| 51 | 35.8 | 37.1 | 36.8 | 36.4 | NA |
| 52 | NA | NA | NA | NA | NA |
| 53 | NA | NA | NA | NA | NA |
| 54 | 20.8 | 22.6 | 22.7 | 17.9 | NA |
| 55 | 0.7 | NA | NA | NA | NA |
| 56 | NA | NA | NA | NA | NA |
| 57 | NA | 0.5 | NA | 0.5 | NA |
| 58 | NA | NA | NA | NA | NA |
| 59 | NA | NA | NA | NA | NA |
| 60 | NA | NA | NA | NA | NA |
| 61 | NA | NA | NA | NA | NA |
| 62 | NA | NA | NA | NA | NA |
| 63 | NA | NA | NA | NA | NA |
| 64 | NA | NA | NA | NA | NA |
| 65 | NA | NA | NA | NA | NA |
| 66 | NA | NA | NA | NA | NA |
| 67 | NA | NA | NA | NA | NA |
| 68 | NA | NA | NA | NA | NA |
| 69 | NA | NA | NA | NA | NA |
| 70 | NA | NA | NA | NA | NA |
| 71 | NA | NA | NA | NA | NA |
| 72 | NA | NA | NA | NA | NA |
| 73 | NA | NA | NA | NA | 1 |
| 74 | NA | NA | NA | NA | NA |
| 75 | NA | NA | NA | NA | NA |
| 76 | NA | NA | NA | NA | NA |
| 77 | NA | NA | NA | NA | NA |
| 78 | NA | NA | NA | NA | NA |
| 79 | NA | NA | NA | NA | NA |
| 80 | NA | NA | NA | NA | NA |
| 81 | NA | NA | NA | NA | NA |
| 82 | NA | NA | NA | NA | NA |
| 83 | NA | NA | NA | NA | NA |
| 84 | NA | NA | NA | NA | NA |
| 85 | NA | NA | NA | NA | NA |
| 86 | NA | NA | NA | NA | NA |
| 87 | NA | NA | NA | NA | NA |
| 88 | NA | NA | NA | NA | NA |
| 89 | NA | NA | NA | NA | NA |
| 90 | NA | NA | NA | NA | NA |
| 91 | NA | NA | NA | NA | NA |
| Total | 44.3 | 45.5 | 41.6 | 45.2 | 20.3 |

**Table S6.** Percentage of *Chrysiptera cyanea* contig 1 aligned with *Amphiprion clarkii, A. ocellaris, A. percula, Acanthochromis polyacanthus,* and *Dascyllus trimaculatus* chromosome-scale genomes based on MUMmer 3.23. Alignments shorter than 10,000 bases were dropped.

| Genome | Chromosome | % contig 1 |
| --- | --- | --- |
| *A. clarkii* | 10 | 25.6 |
| *A. clarkii* | 16 | 30.1 |
| *A. ocellaris* | 11 | 25.1 |
| *A. ocellaris* | 16 | 31.6 |
| *Ac. polyacanthus* | 14 | 24.7 |
| *Ac. polyacanthus* | 15 | 28.9 |
| *A. percula* | 12 | 25.1 |
| *A. percula* | 14 | 3.5 |
| *A. percula* | 20 | 27.9 |
| *D. trimaculatus* | 12 | 12.2 |
| *D. trimaculatus* | 20 | 14.1 |

**Figure S1.**
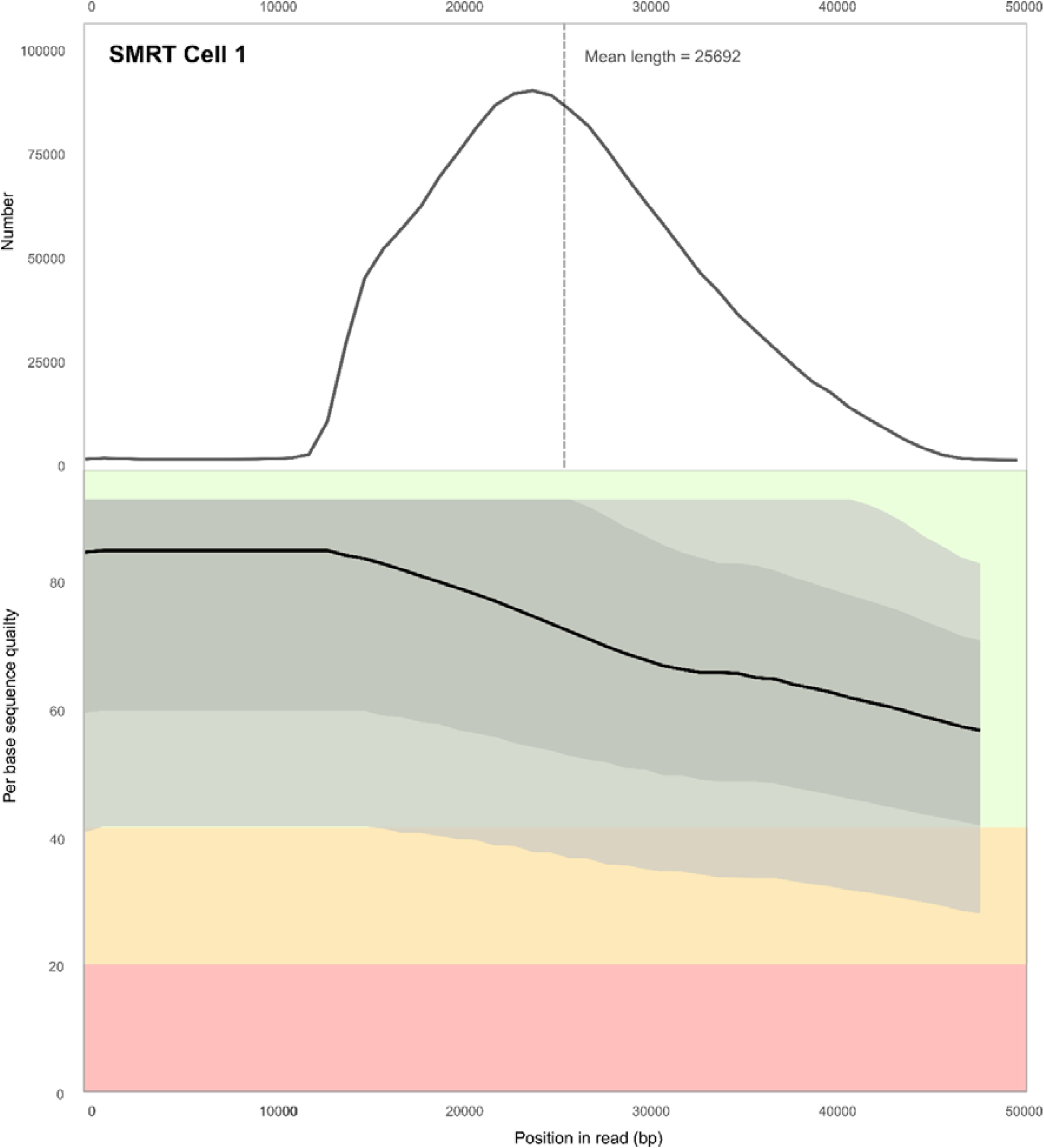
Mean read length and per base sequence quality of the raw reads from the first SMRT cell used for sequencing; data obtained using FastQC 0.11.9.

**Figure S2.**
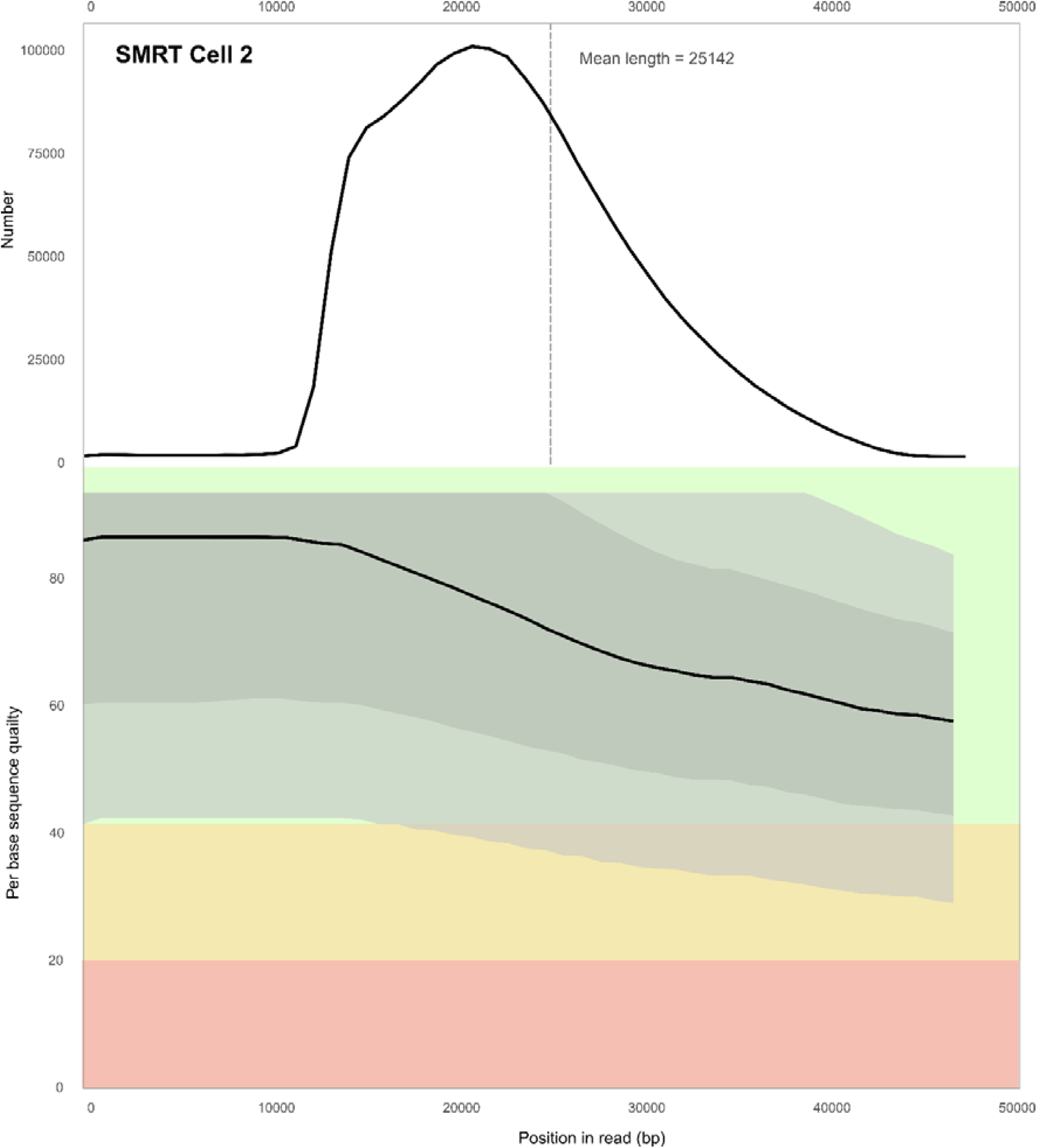
Mean read length and per base sequence quality of the raw reads from the second SMRT cell used for sequencing; data obtained using FastQC 0.11.9.

**Figure S3.**
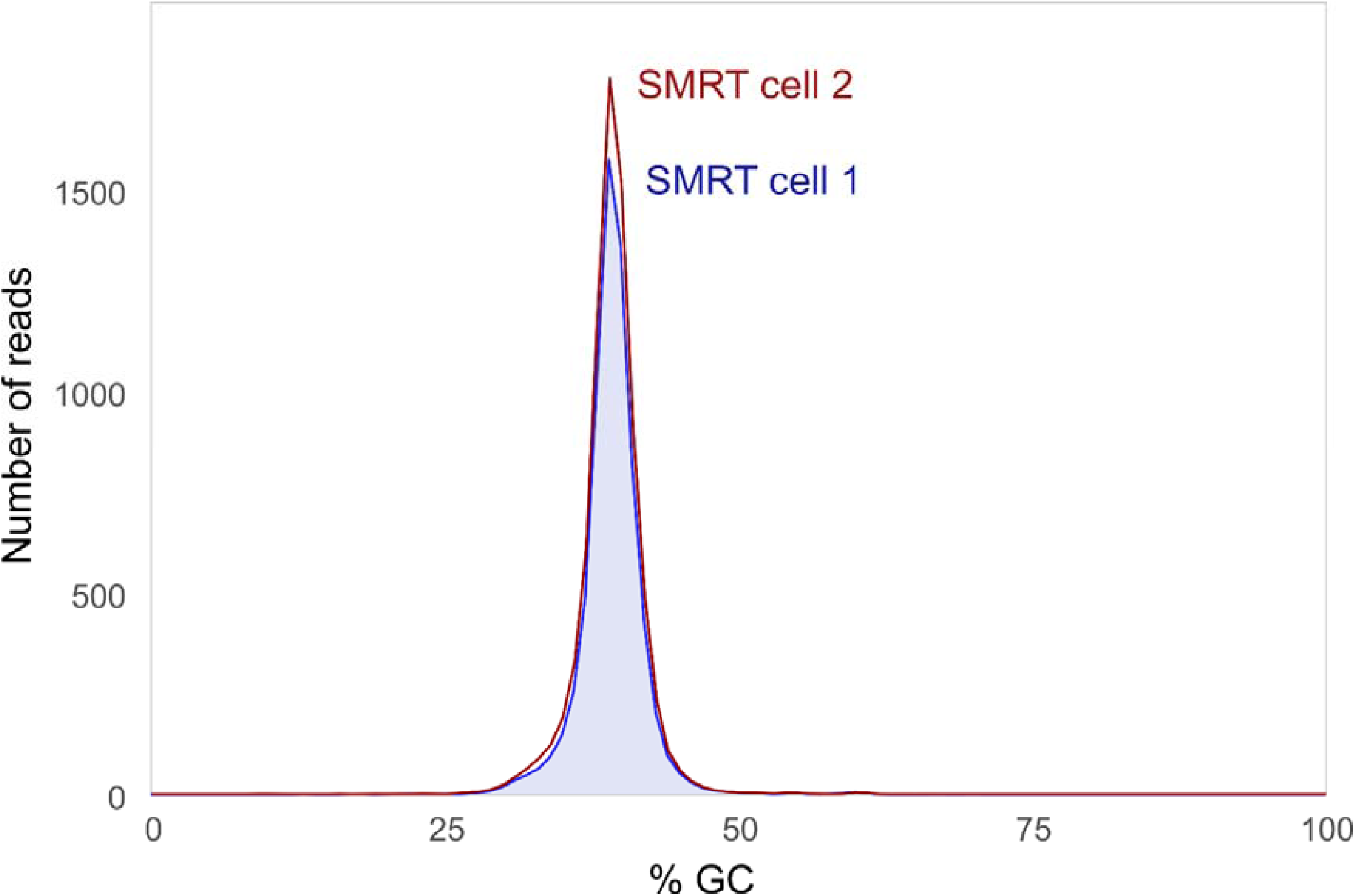
GC content per sequence of the raw reads from the two SMRT cells used for sequencing; data obtained using FastQC 0.11.9.

**Figure S4.**
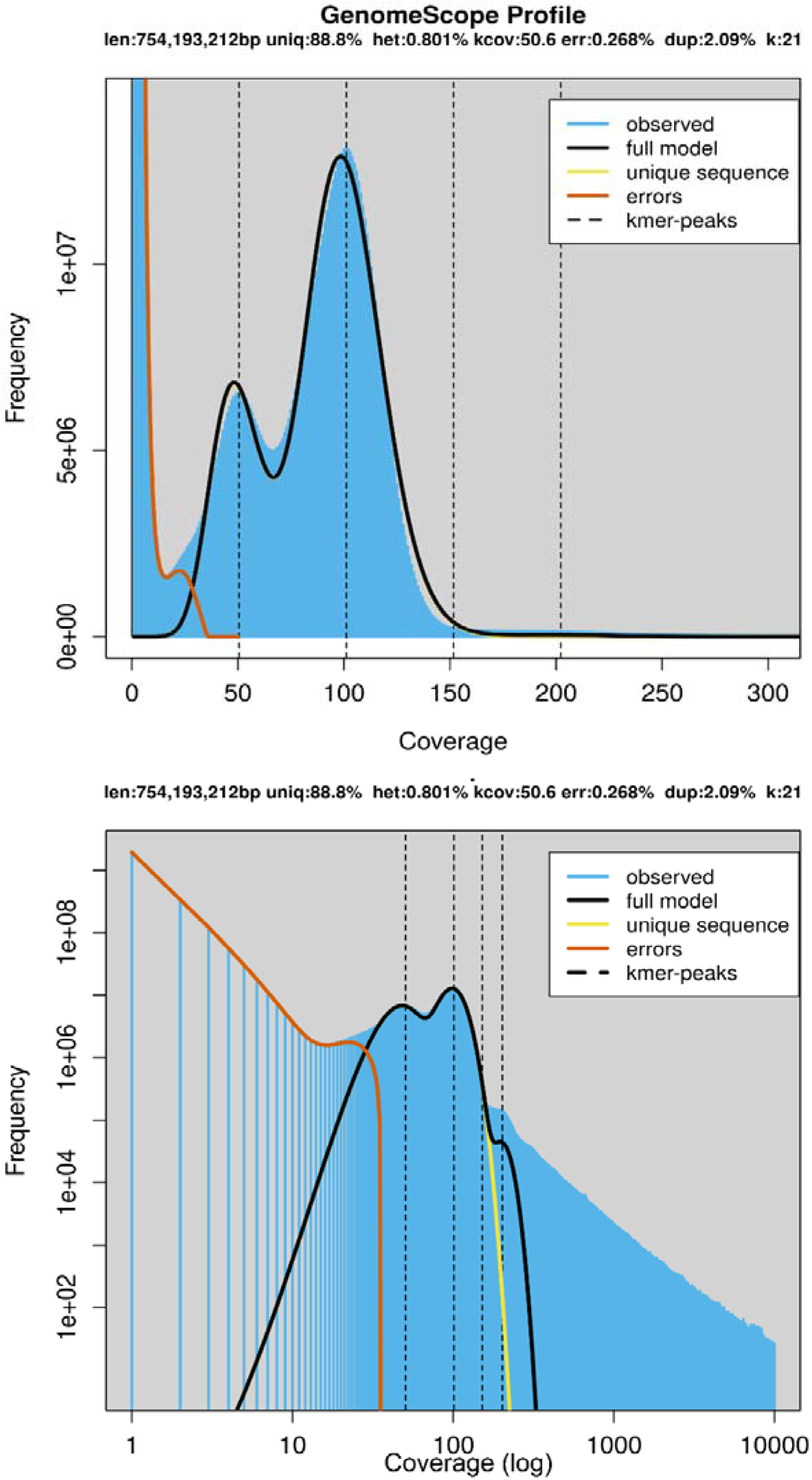
GenomeScope profile of the frequency at which k-mers are covered (raw numbers and log-transformed respectively) within the raw reads, following k-mer assessment by Jellyfish (k-mer size = 21 bases). Figure generated by GenomeScope.

**Figure S4.**
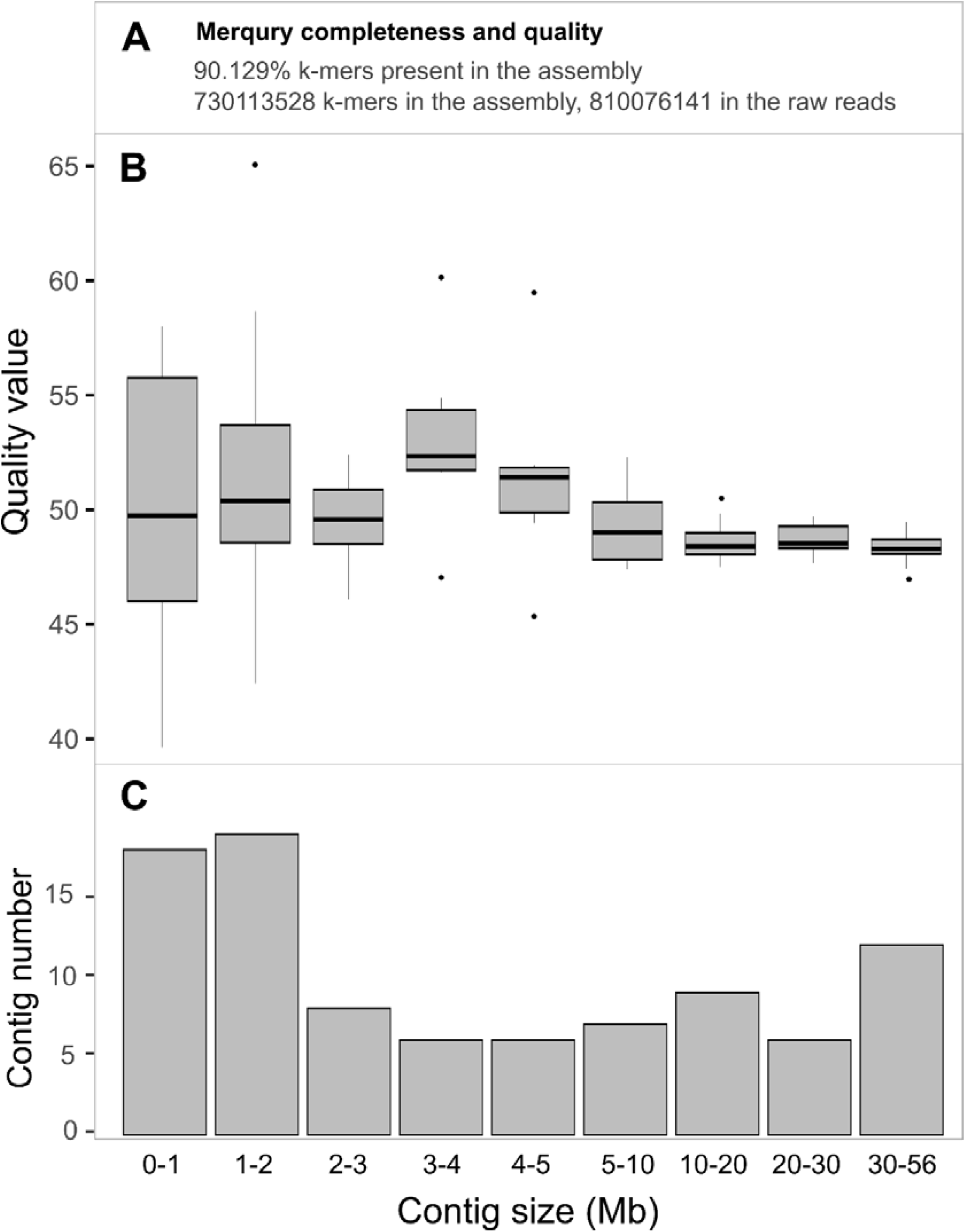
Assembly completeness assessment based on k-mers in the assembly and the raw reads. A: key statistics about the final assembly (IPA no phase, purged). B: Boxplot of the quality value of the contigs based on their size. The boxplot displays the median, 25^th^ and 75^th^ percentiles, with whiskers extending to the maximum and minimum values, within a distance from the box up to 1.5 times the interquartile range. C: Distribution of the contigs in the assembly based on their size. Data obtained using Mercury.

## References

[1] Lehmann R, Lightfoot DJ, Schunter C, Michell CT, Ohyanagi H, Mineta K, et al. Finding Nemo’s Genes: A chromosome-scale reference assembly of the genome of the orange clownfish Amphiprion percula. Molecular Ecology Resources 2019;19:570–85. 10.1111/1755-0998.12939.

[2] Marcionetti A, Rossier V, Roux N, Salis P, Laudet V, Salamin N. Insights into the Genomics of Clownfish Adaptive Radiation: Genetic Basis of the Mutualism with Sea Anemones. Genome Biology and Evolution 2019;11:869–82. 10.1093/gbe/evz042.

[3] Roberts MB, Schultz DT, Gatins R, Escalona M, Bernardi G. Chromosome-level genome of the three-spot damselfish, Dascyllus trimaculatus. G3 (Bethesda) 2023;13:jkac339. 10.1093/g3journal/jkac339.

[4] Ryu T, Herrera M, Moore B, Izumiyama M, Kawai E, Laudet V, et al. A chromosome-scale genome assembly of the false clownfish, Amphiprion ocellaris. G3 Genes|Genomes|Genetics 2022;12:jkac074. 10.1093/g3journal/jkac074.

[5] Schunter C, Welch MJ, Ryu T, Zhang H, Berumen ML, Nilsson GE, et al. Molecular signatures of transgenerational response to ocean acidification in a species of reef fish. Nature Clim Change 2016;6:1014–8. 10.1038/nclimate3087.

[6] Tan MH, Austin CM, Hammer MP, Lee YP, Croft LJ, Gan HM. Finding Nemo: hybrid assembly with Oxford Nanopore and Illumina reads greatly improves the clownfish (Amphiprion ocellaris) genome assembly. GigaScience 2018;7:gix137. 10.1093/gigascience/gix137.

[7] Sayers EW, Bolton EE, Brister JR, Canese K, Chan J, Comeau DC, et al. Database resources of the national center for biotechnology information. Nucleic Acids Res 2022;50:D20–6. 10.1093/nar/gkab1112.

[8] Lehmann R, Schunter C, Welch MJ, Arold ST, Nilsson GE, Tegner JN, et al. Genetic architecture of behavioural resilience to ocean acidification 2022:2022.10.18.512656. 10.1101/2022.10.18.512656.

[9] Moore B, Herrera M, Gairin E, Li C, Miura S, Jolly J, et al. The chromosome-scale genome assembly of the yellowtail clownfish Amphiprion clarkii provides insights into melanic pigmentation of anemonefish 2022:2022.07.21.500941. 10.1101/2022.07.21.500941.

[10] Quoy J, Gaimard J. Zoologie. Voyage autour du monde, exécuté sur les corvettes de S.M. l’Uranie et la Physicienne pendant les années 1817, 1818, 1819 et 1820. Pillet Aîné, Paris: Freycinet L; 1825, p. 392.

[11] Allen GR. Damselfishes of the world. Melle, Germany, Mentor, Ohio: Mergus□; Aquarium Systems [distributor]; 1991.

[12] Gronell AM. Visiting Behaviour by Females of the Sexually Dichromatic Damselfish, Chrysiptera cyanea (Teleostei: Pomacentridae): a Probable Method of Assessing Male Quality. Ethology 1989;81:89–122. 10.1111/j.1439-0310.1989.tb00760.x.

[13] Tamilmani G, Gopakumar G. Chrysiptera cyanea (Quoy & Gaimard, 1825), Kochi: ICAR – Central Marine Fisheries Research Institute; 2017, p. 301–5.

[14] Wacker S, Ness MH, Östlund-Nilsson S, Amundsen T. Social structure affects mating competition in a damselfish. Coral Reefs 2017;36:1279–89. 10.1007/s00338-017-1623-4.

[15] Bapary MAJ, Fainuulelei P, Takemura A. Environmental control of gonadal development in the tropical damselfish Chrysiptera cyanea. Marine Biology Research 2009;5:462–9. 10.1080/17451000802644722.

[16] Bapary JMA, Nurul Amin Md, Takemura A. Food availability as a possible determinant for initiation and termination of reproductive activity in the tropical damselfish Chrysiptera cyanea. Marine Biology Research 2012;8:154–62. 10.1080/17451000.2011.605146.

[17] Gopakumar G, Santhosi I, Ramamoorthy N. Breeding and larviculture of the sapphire devil damselfish Chrysiptera cyanea. Journal of the Marine Biological Association of India 2009;51:130–6.

[18] Thresher RE, Moyer JT. Male success, courtship complexity and patterns of sexual selection in three congeneric species of sexually monochromatic and dichromatic damselfishes (Pisces: Pomacentridae). Animal Behaviour 1983;31:113–27. 10.1016/S0003-3472(83)80179-1.

[19] Steinke D, Zemlak TS, Hebert PDN. Barcoding Nemo: DNA-Based Identifications for the Ornamental Fish Trade. PLOS ONE 2009;4:e6300. 10.1371/journal.pone.0006300.

[20] Nakabo T. Fishes of Japan with Pictorial Keys to the Species [Japanese]. 3rd ed. Tokai University Press; 2013.

[21] Lee S-G, Na D, Park C. Comparability of reference-based and reference-free transcriptome analysis approaches at the gene expression level. BMC Bioinformatics 2021;22:310. 10.1186/s12859-021-04226-0.

[22] Herrera M, Ravasi T, Laudet V. Anemonefishes: A model system for evolutionary genomics. F1000Res 2023;12:204. 10.12688/f1000research.130752.2.

[24] Sovic I, Kronenberg Z, Dunn C, Barnett D, Kingan S, Drake J. Improved Phased Assembler – PacBio HiFi genome assembly 2020.

[24] Kolmogorov M, Yuan J, Lin Y, Pevzner P. Assembly of Long Error-Prone Reads Using Repeat Graphs. Nature Biotechnology 2019;37:540–6. doi:10.1038/s41587-019-0072-8.

[25] Gurevich A, Saveliev V, Vyahhi N, Tesler G. QUAST: quality assessment tool for genome assemblies. Bioinformatics 2013;29:1072–5. 10.1093/bioinformatics/btt086.

[26] Seppey M, Manni M, Zdobnov EM. BUSCO: Assessing Genome Assembly and Annotation Completeness. In: Kollmar M, editor. Gene Prediction: Methods and Protocols, New York, NY: Springer; 2019, p. 227–45. 10.1007/978-1-4939-9173-0_14.

[27] Roach MJ, Schmidt SA, Borneman AR. Purge Haplotigs: allelic contig reassignment for third-gen diploid genome assemblies. BMC Bioinformatics 2018;19:460. 10.1186/s12859-018-2485-7.

[28] Flynn JM, Hubley R, Goubert C, Rosen J, Clark AG, Feschotte C, et al. RepeatModeler2 for automated genomic discovery of transposable element families. Proceedings of the National Academy of Sciences 2020;117:9451–7. 10.1073/pnas.1921046117.

[29] Tempel S. Using and understanding RepeatMasker. Methods Mol Biol 2012;859:29–51. 10.1007/978-1-61779-603-6_2.

[30] Quinlan AR, Hall IM. BEDTools: a flexible suite of utilities for comparing genomic features. Bioinformatics 2010;26:841–2. 10.1093/bioinformatics/btq033.

[31] Kurtz S, Phillippy A, Delcher AL, Smoot M, Shumway M, Antonescu C, et al. Versatile and open software for comparing large genomes. Genome Biology 2004.

[32] Wickham H. ggplot2: Elegant Graphics for Data Analysis 2016.

[33] Monlong J. MUMmerplots with ggplot2, Hippocamplus 2017.

[34] Andrews S, Biggins L, Inglesfield S, Montgomery J. Babraham Bioinformatics – FastQC A Quality Control tool for High Throughput Sequence Data. FastQC A Quality Control Tool for High Throughput Sequence Data 2019. https://www.bioinformatics.babraham.ac.uk/projects/fastqc/ (accessed March 1, 2023).

[35] Bolger AM, Lohse M, Usadel B. Trimmomatic: a flexible trimmer for Illumina sequence data. Bioinformatics 2014;30:2114–20. 10.1093/bioinformatics/btu170.

[36] Kim D, Langmead B, Salzberg SL. HISAT: a fast spliced aligner with low memory requirements. Nat Methods 2015;12:357–60. 10.1038/nmeth.3317.

[37] Li H, Handsaker B, Wysoker A, Fennell T, Ruan J, Homer N, et al. The Sequence Alignment/Map format and SAMtools. Bioinformatics 2009;25:2078–9. 10.1093/bioinformatics/btp352.

[38] Bray NL, Pimentel H, Melsted P, Pachter L. Near-optimal probabilistic RNA-seq quantification. Nat Biotechnol 2016;34:525–7. 10.1038/nbt.3519.

[39] Barnett DW, Garrison EK, Quinlan AR, Strömberg MP, Marth GT. BamTools: a C++ API and toolkit for analyzing and managing BAM files. Bioinformatics 2011;27:1691–2. 10.1093/bioinformatics/btr174.

[40] Bruna T, Hoff KJ, Lomsadze A, Stanke M, Borodovsky M. BRAKER2: Automatic Eukaryotic Genome Annotation with GeneMark-EP+ and AUGUSTUS Supported by a Protein Database. NAR Genomics and Bioinformatics 2021;3:lqaa108.

[41] Bruna T, Lomsadze A, Borodovsky M. GeneMark-EP+: eukaryotic gene prediction with self-training in the space of genes and proteins. NAR Genomics and Bioinformatics 2020;2:lqaa026.

[42] Buchfink B, Reuter K, Drost H-G. Sensitive protein alignments at tree-of-life scale using DIAMOND. Nat Methods 2021;18:366–8. 10.1038/s41592-021-01101-x.

[43] Gotoh O. A space-efficient and accurate method for mapping and aligning cDNA sequences onto genomic sequence. Nucleic Acids Research 2008;36:2630–8.

[44] Hoff KJ, Lomsadze A, Borodovsky M, Stanke M. Whole-genome annotation with BRAKER. Gene Prediction, New York, NY: Humana; 2019, p. 65–95.

[45] Hoff KJ, Lange S, Lomsadze A, Borodovsky M, Stanke M. BRAKER1: unsupervised RNA-Seq-based genome annotation with GeneMark-ET and AUGUSTUS. Bioinformatics 2016;32:767–9.

[46] Iwata H, Gotoh O. Benchmarking spliced alignment programs including Spaln2, an extended version of Spaln that incorporates additional species-specific features. Nucleic Acids Research 2012;40:e161.

[47] Lomsadze A, Burns PD, Borodovsky M. Integration of mapped RNA-Seq reads into automatic training of eukaryotic gene finding algorithm. Nucleic Acids Res 2014;42:e119. 10.1093/nar/gku557.

[48] Lomsadze A, Ter-Hovhannisyan V, Chernoff, YO, Borodovsky M. Gene identification in novel eukaryotic genomes by self-training algorithm. Nucleic Acids Research 2005;33:6494–506.

[49] Stanke M, Diekhans M, Baertsch R, Haussler D. Using native and syntenically mapped cDNA alignments to improve de novo gene finding. Bioinformatics 2008;24:637–44. 10.1093/bioinformatics/btn013.

[50] Stanke M, Schöffmann O, Morgenstern B, Waack S. Gene prediction in eukaryotes with a generalized hidden Markov model that uses hints from external sources. BMC Bioinformatics 2006;7:62. 10.1186/1471-2105-7-62.

[51] UniProt Consortium. UniProt: the universal protein knowledgebase in 2021. Nucleic Acids Res 2021;49:D480–9. 10.1093/nar/gkaa1100.

[52] Zdobnov EM, Apweiler R. InterProScan – an integration platform for the signature-recognition methods in InterPro. Bioinformatics 2001;17:847–8. 10.1093/bioinformatics/17.9.847.

[53] Altschul SF, Madden TL, Schäffer AA, Zhang J, Zhang Z, Miller W, et al. Gapped BLAST and PSI-BLAST: a new generation of protein database search programs. Nucleic Acids Res 1997;25:3389–402. 10.1093/nar/25.17.3389.

[54] Condon K. tispec: Calculates Tissue Specificity from RNA-Seq Data 2020.

[55] Conway JR, Lex A, Gehlenborg N. UpSetR: an R package for the visualization of intersecting sets and their properties. Bioinformatics 2017;33:2938–40. 10.1093/bioinformatics/btx364.

[56] Moore B, Herrera M, Gairin E, Li C, Miura S, Jolly J, et al. The chromosome-scale genome assembly of the yellowtail clownfish Amphiprion clarkii provides insights into the melanic pigmentation of anemonefish. G3 Genes|Genomes|Genetics 2023;13:jkad002. 10.1093/g3journal/jkad002.

[57] Kryuchkova-Mostacci N, Robinson-Rechavi M. A benchmark of gene expression tissue-specificity metrics. Briefings in Bioinformatics 2017;18:205–14. 10.1093/bib/bbw008.

[58] Yanai I, Benjamin H, Shmoish M, Chalifa-Caspi V, Shklar M, Ophir R, et al. Genome-wide midrange transcription profiles reveal expression level relationships in human tissue specification. Bioinformatics 2005;21:650–9. 10.1093/bioinformatics/bti042.

[59] Lecchini D, Adjeroud M, Pratchett MS, Cadoret L, Galzin R. Spatial structure of coral reef fish communities in the Ryukyu Islands, southern Japan. Oceanologica Acta 2003;26:537–47. 10.1016/S0399-1784(03)00048-3.

[60] McCord CL, Nash CM, Cooper WJ, Westneat MW. Phylogeny of the damselfishes (Pomacentridae) and patterns of asymmetrical diversification in body size and feeding ecology. PLOS ONE 2021;16:e0258889. 10.1371/journal.pone.0258889.

[61] Tang KL, Stiassny MLJ, Mayden RL, DeSalle R. Systematics of Damselfishes. Cope1 2021;109:258–318. 10.1643/i2020105.

[62] Kass JM, Vilela B, Aiello-Lammens ME, Muscarella R, Merow C, Anderson RP. Wallace: A flexible platform for reproducible modeling of species niches and distributions built for community expansion. Methods in Ecology and Evolution 2018;9:1151–6. 10.1111/2041-210X.12945.

[63] QGIS Association. QGIS Geographic Information System 3.22.9 2022.

